# A lineage-resolved molecular atlas of *C. elegans* embryogenesis at single cell resolution

**DOI:** 10.1101/565549

**Authors:** Jonathan S. Packer, Qin Zhu, Chau Huynh, Priya Sivaramakrishnan, Elicia Preston, Hannah Dueck, Derek Stefanik, Kai Tan, Cole Trapnell, Junhyong Kim, Robert H. Waterston, John I. Murray

## Abstract

*C. elegans* is an animal with few cells, but a striking diversity of cell types. Here, we characterize the molecular basis for their specification by profiling the transcriptomes of 84,625 single embryonic cells. We identify 284 terminal and pre-terminal cell types, mapping most single cell transcriptomes to their exact position in *C. elegans’* invariant lineage. We use these annotations to perform the first quantitative analysis of the relationship between lineage and the transcriptome for a whole organism. We find that a strong lineage-transcriptome correlation in the early embryo breaks down in the final two cell divisions as cells adopt their terminal fates and that most distinct lineages that produce the same anatomical cell type converge to a homogenous transcriptomic state. Users can explore our data with a graphical application “VisCello”.

---

To understand how cell fates are specified during development, it is essential to know the temporal sequence of gene expression in cells during their trajectories from uncommitted precursors to differentiated terminal cell types. Gene expression patterns near branch points in these trajectories can help identify candidate regulators of cell fate decisions (*1*). Single cell RNA sequencing (sc-RNA-seq) has made it possible to obtain comprehensive measurements of gene expression in whole animals (*2–6*) and embryos (*7–12*). sc-RNA-seq profiling of multiple developmental stages in a time series can be particularly informative, as algorithms can use the data to reconstruct the developmental trajectories followed by specific cell types. However, confounding factors can generate misleading trajectories. For example, progenitor cell populations with distinct lineage origins may be conflated if their transcriptomes are too similar, and abrupt changes in gene expression can result in gapped trajectories. Thus, information from independent assays is necessary to conclusively validate an inferred trajectory as an accurate model of development. To date, no study has both comprehensively reconstructed and validated developmental trajectories for a whole organism.

The nematode worm *Caenorhabditis elegans* develops through a known and invariant cell lineage from the fertilized egg to an adult hermaphrodite with 959 somatic cells (*13, 14*), which creates the potential for a truly comprehensive understanding of its development. Here, we use sc-RNA-seq, the known *C. elegans* lineage, and imaging of fluorescent reporter genes (*15*, *16*) to produce a lineage-resolved single cell atlas of embryonic development that includes trajectories for most individual cells in the organism. We use this dataset to quantitatively analyze the relationship between the cell lineage and the temporal dynamics of gene expression. We find that in the early embryo, lineage distance between cells predicts how different their transcriptomes will be. In the mid-to-late embryo, expression patterns of closely related cells diverge as they adopt their terminal cell fates. This transition largely occurs during a single cell division which, depending on the particular sublineage, produces either a terminal cell or the parent of two terminal cells of related cell type. In many lineages, we observe that transcription factor (TF) expression patterns are selectively inherited in only one of two daughters of a progenitor cell, contributing to their differentiation.

## Single-cell RNA-seq of *C. elegans* embryos

We sequenced the transcriptomes of single cells from *C. elegans* embryos by using the 10x Genomics platform. We assayed loosely synchronized embryos enriched for early cells as well as embryos that had been aged for ∼300, ∼400, and ∼500 minutes after first cleavage. We processed the datasets with a unified pipeline based on the Monocle software package (*17*). After quality control (see **Methods**), the final integrated dataset contained 84,625 single cells, representing >60x oversampling of the 1,341 branches in the *C. elegans* embryonic lineage.

We estimated the embryo stage of each cell by comparing its expression profile with a high-resolution whole-embryo RNA-seq time series (*18*) (**Fig. S1** and **Methods**). We then visualized the data with the Uniform Manifold Approximation and Projection (UMAP) (*19*, *20*) algorithm, which projects the data in low-dimensional space and is well suited for data with complex branching structures (*20*). We found that trajectories in the UMAP projection reflect a smooth progression of embryo time (**Fig. 1A**), with cells collected from later time points usually occupying more peripheral positions (**Fig. 1B**) (see **Supplemental Note 1** for a discussion of the term “trajectory”). Unique transcripts per cell, as estimated using Unique Molecular Identifiers (UMIs), decreased with increasing embryo time, consistent with decreasing physical cell size as embryonic cells divide without growth (**Fig. S2**). These observations suggest that UMAP trajectories corresponded to developmental progression and that embryo time estimates are a reasonable proxy for developmental stage for most cells. Approximately 75% of the cells recovered (64,160 cells) were from embryos of 210-510 minutes, corresponding to the ∼190 cell stage to the 3-fold stage of development (**Fig. 1C**); however, cells were also recovered from early embryos (< 210 minutes, 9,051 cells), and late embryos (> 510 minutes, 11,414 cells).

**Fig 1.**
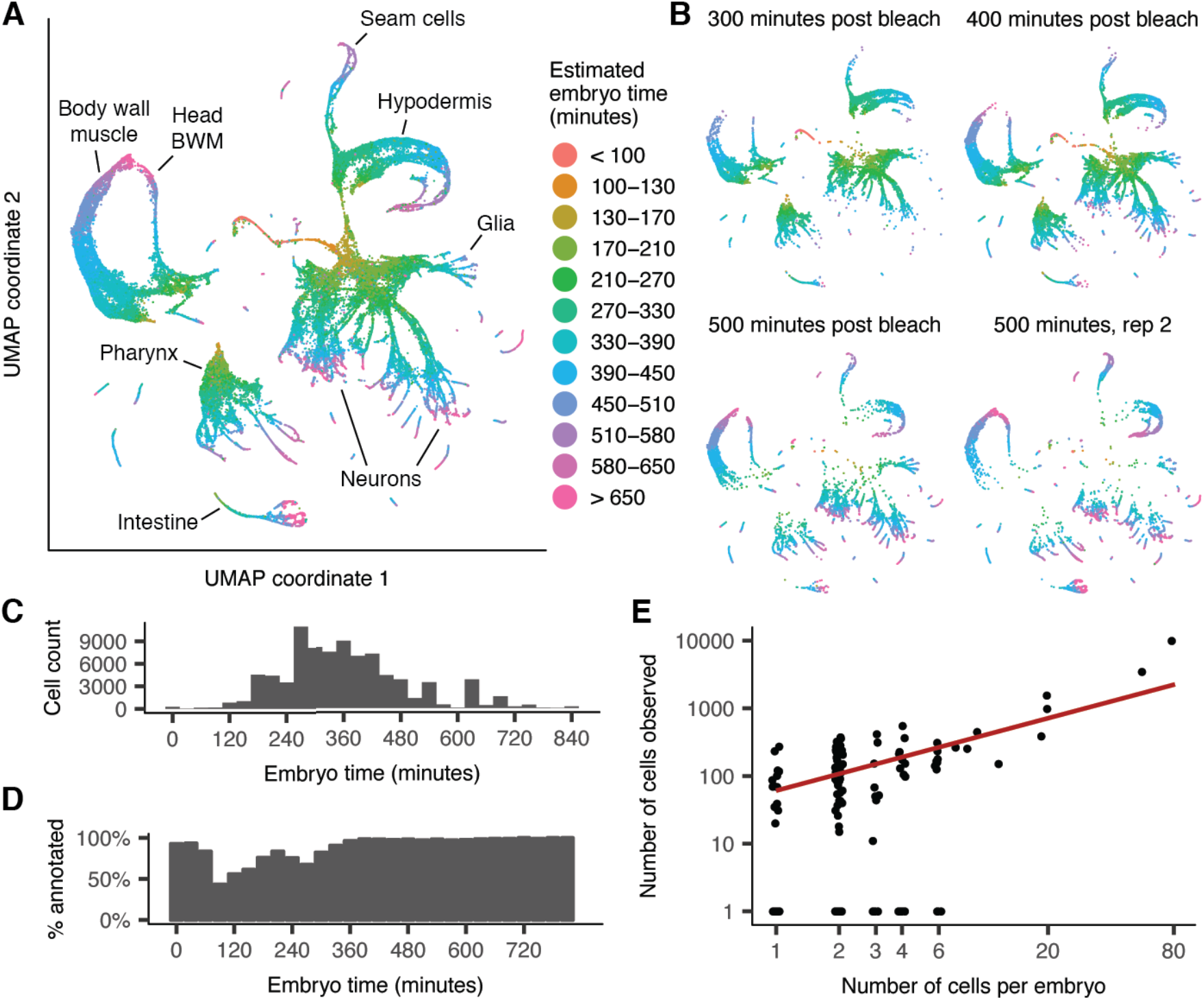
UMAP projection shows tissues and developmental trajectories in C. elegans embryogenesis. (**A**) UMAP projection of the 84,625 cells in the post-QC sc-RNA-seq dataset. Color indicates the age of the embryo that a cell came from, estimated based on correlation to a whole-embryo RNA-seq time series (*18*) and measured in minutes after an embryo’s first cell cleavage. (**B**) Positions of cells from four samples of synchronized embryos (see Methods) on the UMAP plot. (**C**) Histogram of estimated embryo time for all cells in the dataset. (**D**) Bar plot showing for bins of embryo time, the percentage of cells in that embryo time bin for which were able to assign to a terminal cell type or preterminal lineage. (**E**) Scatter plot showing correlation of the number of cells of a given anatomical cell class in a single embryo (X axis, log scale) with the number of cells recovered in our data (Y axis, log scale). Each point corresponds to a cell class. Only cells with embryo time >= 390 minutes are included in the counts (many earlier cells are still dividing). Red line is a linear fit, excluding points with y = 0.

We clustered cells in the UMAP using the Louvain algorithm (*21*) and annotated clusters with cell type identities using marker genes from the extensive literature on *C. elegans* gene expression (*22*) (Markers used for each annotation are listed in **Table S1**). The global UMAP arranges cells into a central group of progenitor cells and branches corresponding to eight major tissues (**Fig. 1A, S3**): muscle/mesoderm, epidermis, pharynx, ciliated neurons, non-ciliated neurons, glia/excretory cells, intestine, and germline. While some individual cell types were identifiable in this global UMAP, many were not, and early progenitor lineages were poorly resolved. To gain resolution, we hierarchically created separate UMAPs of each tissue (**Fig. S4–S11**). These “sub-UMAPs” better resolved specific cell types, allowing us to make extensive, fine-grained annotations.

Using a combination of marker genes, lineage assignments, and developmental time, we located 101 specific terminal anatomical cell types, including every lineage input to body wall muscle, every distinct subtype of pharyngeal muscle (pm1-2, pm3-5, pm6, pm7, and pm8) and hypodermis (hyp1-2, hyp3, hyp4-6, hyp7, hyp8-11, seam, and P cells), and every non-neuronal cell type in the mesoderm. We identified 63 of 82 (75%) non-pharyngeal neuron classes present in the embryo (**Table S2**). A lack of suitable markers limited our ability to identify 12 of 14 pharyngeal neuron classes, and glia other than AMsh, AMso, CEPso, and ILso, despite multiple distinct clusters being evident in the UMAP (**Fig. S9**). We did not identify 7 of 11 tail neuron classes and a few specialized epithelial cell types in the pharynx and rectum, with no obvious unannotated clusters as candidates. Their absence could result from incomplete dissociation of the relevant tissues, damage to the cells causing them to fail QC filtering, or a failure to resolve them from other cell types in the UMAP projections.

Looking across all cells sequenced in our dataset, we successfully annotated 94% of cells from embryo time >300 minutes with a cell type (**Fig. 1D**), and 73% of cells from embryo time < 300 minutes with a cell lineage (discussed below). The number of cells annotated for each cell type was variable but roughly fit the expectation based on the number of cells of that type present in a single embryo (**Fig. 1E**, *r* = 0.64, p = 3.1e-12).

## Annotation of the early lineage

The structure of the global and single-tissue UMAPs was dominated by trajectories of terminal cell differentiation. We hypothesized that closely related lineages could be better resolved by separately analyzing early cells prior to terminal differentiation. Thus, we ran UMAP with only early cells (embryo time <= 150 and <= 250 minutes) and found branching patterns that reflect early lineage identities (**Fig. 2, S12**). Early intestine and germline cells commit to their terminal fates very early and have very divergent expression that distorts the projections, so they were analyzed separately (**Fig. S7, S11**). The 250-minute UMAP contained several large quasi-connected groups corresponding approximately to major founding lineages, roughly organized by the major fates produced by each lineage (MS muscle, MS pharynx, C/D muscle and AB-derived lineages that produce either pharynx, neurons/glia, or hypodermis). We were able to resolve additional details by recursively making sub-UMAP projections of these cell subsets (**Fig. S12, S13**, **Fig. 2D**).

**Fig 2.**
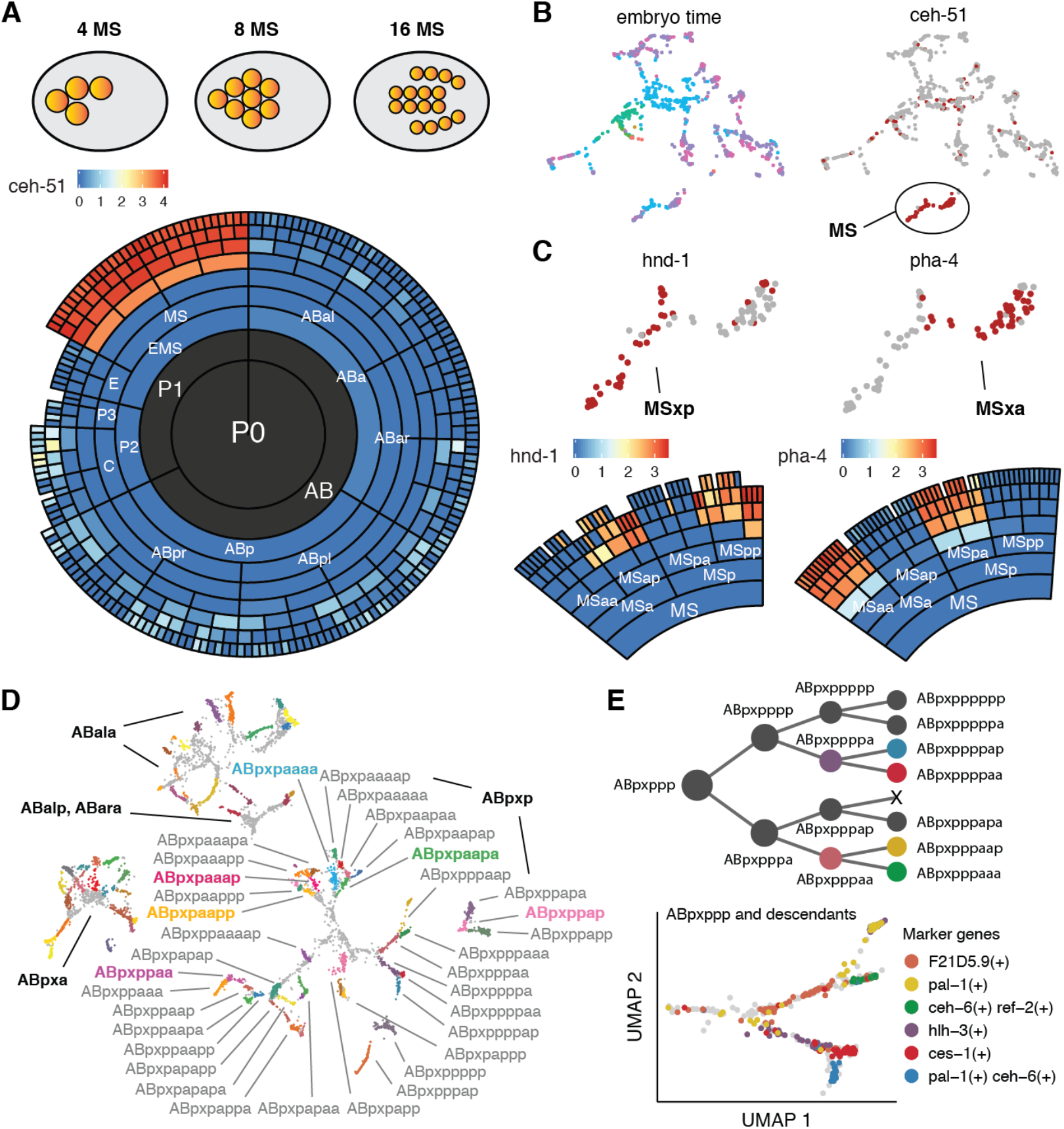
Annotation of the early lineage. (**A**) Diagram showing position of early MS lineage cells marked by expression of ceh-51. The radiograph shows the average fluorescent intensity (log10 scaled) of ceh-51 measured by live imaging. (**B**) UMAP projection of early-stage cells (estimated embryo time <= 150 minutes) colored by time and ceh-51 expression. (**C**) Expression of hnd-1 and pha-4 measured by sc-RNA-seq (UMAP) and live imaging (radiograph). (**D**) Annotation of AB lineage early stage cells (estimated embryo time <= 250 minutes) that give rise to neurons and glia, with full annotation of the ABpxp lineage. Colored bold annotations highlight specific lineages that are discussed in the text. Full annotation of this UMAP is shown in **Fig. S11**. (**E**) Lineage tree of for the ABpxppp sub-lineage, highlighting cells that are present in (D). The (co-)expression pattern of marker genes identifies branches in the UMAP that correspond to specific ABpxppp descendants.

To annotate these early lineages, we exploited the EPiC database, which contains single cell resolution expression profiles extracted by cell tracking software from confocal movies of *C. elegans* embryos expressing fluorescent reporters (*15*). In addition to the 180 previously described patterns (*15*, *23*), we collected movies for 71 additional genes, increasing the total number of patterns in EPiC to 251 genes (**Table S3**). We annotated branches with lineage identities between the 50-cell and 350-cell stages by finding genes that were differentially expressed both between sister lineages in the EPiC data and between branches of the sub-UMAP trajectories in a concordant manner (**Fig. 2**, **Fig. S12**, **Tables S4-S5**). For example, expression of *ceh-51* is restricted to the MS (mesoderm-producing) lineage (*24*), allowing us to label the single group of *ceh-51(+*) cells in 150-minute UMAP as the MS lineage (**Fig. 2A, 2B**). Within this lineage, we used expression of *pha-4* to annotate the anterior granddaughters of MS (MSaa and MSpa) and *hnd-1* to annotate the posterior granddaughters (MSap and MSpp) (**Fig. 2C**). We applied this same logic iteratively across the different UMAPs and lineage marker genes to annotate each branch with its lineage identity. Markers used to annotated each lineage are described in **Table S4**.

In most cases, branches in the early embryo UMAPs corresponded directly to sister cells in the lineage (**Fig. 2D, 2E**), but some branches were unclear or misleading, and marker gene expression was critical to identify the correct annotations. For example, ABpxpaaaa and ABpxpaapa are cousin lineages, but appear to branch as sisters in the UMAP trajectory, and the same is true for their sisters (ABpxpaaap and ABpxpaapp) (**Fig. 2D**). In other cases, such as the ABpxppap lineage (**Fig. 2D**), marker gene combinations were required to annotate lineages that were not contiguous with their parent or sister lineages in the UMAP. These misleading branches demonstrate the importance of having independent expression or lineage data to correctly interpret estimated trajectories visualized in low-dimensional embeddings of sc-RNA-seq data.

In total, we annotated 183 pre-terminal lineages. Our annotations were most comprehensive for the 350-cell stage (∼160-210 minutes), in which we identified 79% of AB lineage cells, 88% of MS lineage cells, and all C lineage cells (cells from the clonal D and E lineages were also annotated, but were too homogenous to narrow down to exact 350-cell stage cells). These annotations included immediate progenitors of several terminal cell types that we could not directly identify. In total, 80.0% of terminal anatomical cells in the embryo were directly annotated, and an additional 16.4% had at least a parent or grandparent annotated, leaving only 3.6% unrepresented. Overall, our annotations represent a substantially larger fraction of the full cell lineage of an organism, at single-lineage resolution, than any previous large-scale sc-RNA-seq study of development (*7–12*).

The aggregate gene expression profiles for each terminal cell type (binned by embryo time) and each pre-terminal cell lineage are reported in **Tables S6 and S7** respectively.

## Cell fate decisions of ciliated neurons

Developmental trajectories in which a parent cell divides to produce two terminal daughter cells of different cell types are a basic type of cell fate decision. Bifurcations like these are common in neuronal lineages, such as those that produce ciliated neurons. To examine the molecular basis for such developmental decisions, we used a recursive UMAP projection of ciliated neurons to identify developmental trajectories for 20 of 22 ciliated neuron types (**Fig. 3A**), all of which included both the terminal neuron and its parent. The progenitors of the two unidentified neurons (PHA and PHB), were located in a sub-clustering of non-ciliated neurons, consistent with the fact that these neurons’ sister cells produce non-ciliated neurons (**Fig. S10**). The distinction between neuroblasts and terminal neurons was supported by embryo time estimates consistent with terminal cell division times (*25*), by the expression patterns of cell cycle associated genes and transcription factors (**Fig. 3B**), and by the structure of the UMAP projection. A 3D version of the UMAP featured better continuity for several trajectories, including those connecting the ASG-AWA, ADF-AWB, and ASJ-AUA neuroblasts with their daughter cells, as well as the branching of the laterally asymmetric left and right ASE neurons (**Fig. S14**).

**Fig 3.**
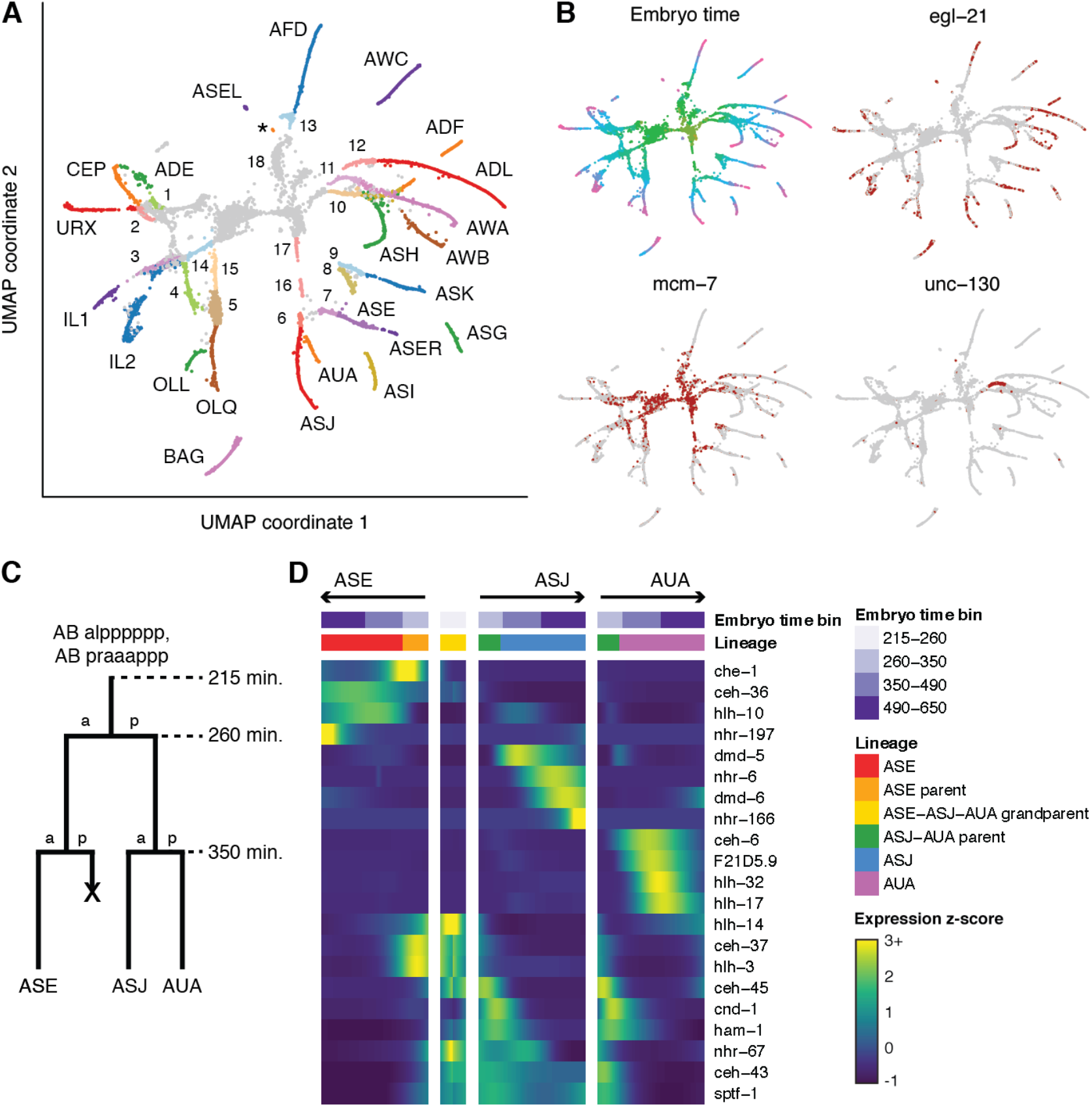
Developmental trajectories of ciliated neurons. (**A**) UMAP of ciliated neurons and precursors. Colors correspond to cell identity. Text labels indicate terminal cells. Numbers 1-13 indicate parents of **1** ADE-ADA, **2** CEP-URX **3** IL1 **4** OLL **5** OLQ **6** ASJ-AUA **7** ASE **8** ASI **9** ASK **10** ADF-AWB **11** ASG-AWA **12** ADL **13** AFD-RMD. **3-5**, **7-9**, and **12** are listed as parents of only one cell type as the sister cells die. Numbers 14-17 indicate grandparents of **14** IL1 (= IL2 parent) **15** OLQ-URY **16**, **17** ASE-ASJ-AUA. **18** indicates a progenitor cluster that includes the AWC-SAAVx and BAG-SMDVx parents, which were identified in a separate UMAP (**Fig. S12C**). This latter analysis also tentatively identified a few cells near the base of the ASH trajectory as the ASH-RIB parent. Late stage AUA cells cluster with non-ciliated neurons and are not included in this UMAP but are included in the heatmap in panel D. The tiny cluster of cells labeled with an asterisk (*) is putatively AWC-ON based on srt-28 expression. (**B**) UMAP plot colored by embryo time (colors matched to **Fig. 1A**) and gene expression (red indicates >0 reads for the listed gene). mcm-7 is gene associated with the cell cycle. unc-130 is known to be expressed in the ASG-AWA neuroblast but neither terminal cell (*40*) (**C**) Cartoon illustrating the lineage of the ASE, ASJ, and AUA neurons. (**D**) Heatmap showing patterns of differential transcription factor expression associated with branches in the ASE-ASJ-AUA lineage. Expression values are log-transformed, then centered and scaled by standard deviation for each row (gene).

To identify potential regulators of cell fate decisions, we identified genes that were differentially expressed between the branches of each bifurcating ciliated neuron lineage (**Fig. 3C-D, S15**). The lineage of the ASE, ASJ, and AUA neurons (spanning embryo time ∼215-650 minutes) serves as a representative example (**Fig. 3C**). About 3-4 TFs are differentially expressed in each branch of this lineage compared to its sister (**Fig. 3D**). Beyond these simple cases, we also found several TFs that were expressed in a parent cell and selectively retained in one daughter but not the other. For example, the TFs *ceh-36/37/43/45* and *ham-1* are all coexpressed within single ASE-ASJ-AUA neuroblast cells. *ceh-36* and *ceh-37* expression was retained in only one daughter of this neuroblast, the ASE parent, while *ceh-43, ceh-45*, and *ham-1* expression was retained only in the other daughter, the ASJ-AUA neuroblast. Similarly, *ceh-43, sptf-1*, and *nhr-67* are co-expressed in the ASJ-AUA neuroblast and its daughter ASJ, but are lost in AUA.

Other lineage bifurcations that produce two different neurons showed similar patterns of differential gene expression (**Fig. S15**). Neurons (like ASE) with a sister cell that undergoes cell death expressed on average fewer type-specific TFs that distinguish them from both their parent and other terminal neurons, compared to neurons with sisters that are also neurons of a different type (**Fig. S16**, p = 0.032, Wilcoxon rank sum test). Consistent with the hypothesis that these lineages may have already started to commit to a cell fate in the parent’s generation, several of them featured expression of a neuron type-specific TF in the parent of terminal neuron—*che-1* and *ceh-36* for ASE, *ttx-3* for ASK, *sox-3* for OLQ, and *ceh-8* for RIA. This expression suggests that these neuroblasts may begin committing to a cell fate early, before the terminal cell is produced.

Only two neurons (ASE and AWC) are known to have left-right asymmetric gene expression (*28*, *29*). For both neuron types, the lineages of the left and right neurons diverge in the early embryo at the 4-cell stage (< 50 minutes). Asymmetric gene expression in our data, however, emerges much later in embryogenesis. The transcriptomes of ASEL and ASER diverged in our UMAP at ∼650-700 minutes and *lim-6* was expressed specifically in the ASEL branch, consistent with previous studies (*30, 31*). AWC left/right asymmetry occurs stochastically, with one neuron becoming “AWC-ON” and the other becoming “AWC-OFF” (*29*). We identified a small cluster in the UMAP with embryo time >700 minutes as AWC-ON based on *srt-28* expression (**Fig. 3A**) (*32*). AWC-OFF is putatively part of the main AWC trajectory. No evidence of left/right asymmetry was observed in neurons besides ASE and AWC.

## Most co-fated lineages converge to a common transcriptomic state

Many of the terminal cell types we identified consist of pairs of cells that are bilaterally symmetric in the body plan and are produced by bilaterally symmetric lineages. Most of these pairs are not distinguishable by UMAP. However, several cell types with >2-fold symmetry are produced by multiple non-symmetric lineage inputs. For these, cell lineages tended to cluster separately in our early cell UMAPs, while in our late-cell tissue UMAPs, we saw almost no evidence of lineage-related heterogeneity within cell types. This difference suggested that the transcriptomes of these co-fated lineages were converging during differentiation.

One example of apparent molecular convergence of cells from distinct lineages was the IL1-IL2 neuroblasts. The six IL1 and six IL2 neurons are produced by three pairs of neuroblast lineages. Each neuroblast pair produces a pair of bilaterally symmetric IL1 neurons, and likewise a pair of IL2 neurons. A UMAP of IL1/2 neurons and progenitors revealed trajectories for these neuroblasts that converge gradually over their lifespan (**Fig. 4A**). The transcription factor *ast-1* was transiently expressed at extremely high levels (>10,000 TPM) during this process, suggesting that it might play a role in homogenizing the IL1/2 neuroblast transcriptomes (**Fig. 4B**). Correspondingly, expression of genes differentially expressed between the input lineages decreased over time, while expression of genes specific to terminal neurons increased (**Fig. 4C-D**).

**Fig 4.**
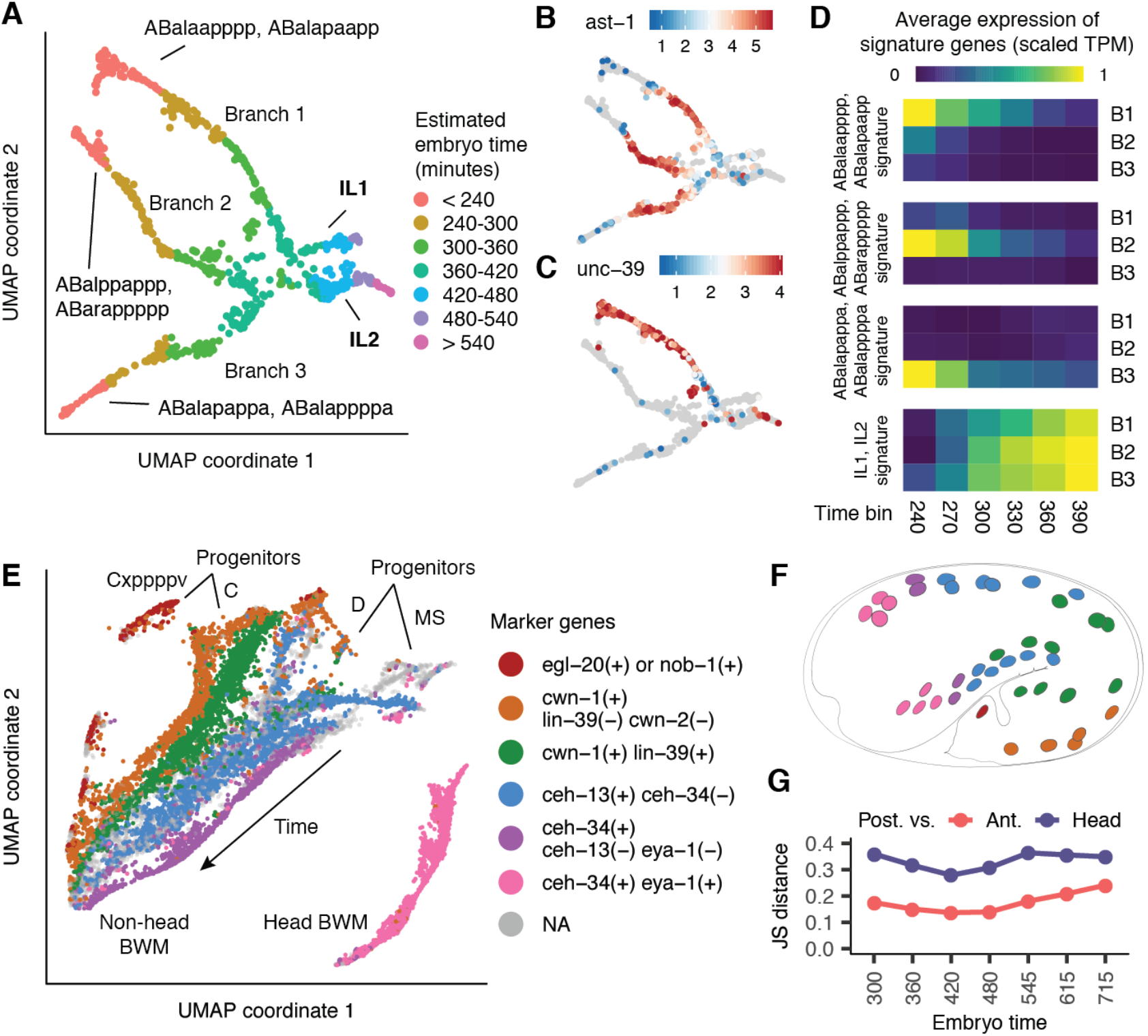
Full vs. incomplete convergence of lineages producing common cell types. (**A**) UMAP of IL1/2 neurons and progenitors colored by estimated embryo time (cells selected based on annotations in **Fig. 3A** and **Fig. S12**). (**B**) IL1/2 UMAP colored by ast-1 expression level (log size-factor normalized UMI counts). (**C**) IL1/2 UMAP colored by expression of unc-39, a gene specific to branch 1. (**D**) Heatmap of lineage and fate signature gene expression across time for each of the 3 branches (see text and **Methods**). (**E**) **Fig. S5A** shows a UMAP of body wall muscle and mesoderm cells. This panel is a zoomed-in view of that UMAP, including only BWM cells, which are grouped into “bands” based on marker gene expression patterns (here, a cell with 0 reads for a gene is considered to still express that gene if >= 2 of 5 of its nearest neighbors have >0 reads). (**F**) Physical positions of cells in each BWM band (colors matched to panel E) in the embryo at 430 minutes. Adapted from **Fig. 8B** of (14). (**G**) Transcriptome JS distance for posterior (orange+green bands in panel E) vs. anterior (blue band) or head (pink band) BWM cells over time. BWM subsets differentiate in parallel, but heterogeneity persists throughout development.

We observed similar lineage convergence via continuous gene expression trajectories for hypodermis (**Fig. S8**), head body wall muscle (**Fig. S13**), and GLR cells (**Fig. S13**). In each of these lineages, the parent of the terminal cells is already committed to the cell fate and does not produce daughters of another cell type. By contrast, other lineages commit to a cell fate only in the final cell division. For example, parents of ILso glia also produce neurons and hypodermis. ILso parents clustered separately from terminal ILso and from each other in our UMAP (**Fig. S17A**). We did not see any intermediate states connecting the parents to the terminal cells, suggesting that the convergence of ILso input lineages may be abrupt. Lineages producing SMD neurons showed similar discontinuities (**Fig. S17B**). Overall, our observations of both continuous and gapped trajectories suggest that different lineages feature different modes of differentiation, with lineages that commit to a cell fate early tending to differentiate gradually, while lineages that commit to a cell fate late differentiate abruptly.

Non-head body wall muscle (BWM) was exceptional in that lineage-related heterogeneity persisted throughout differentiation. BWM is produced by multiple distinct lineages (C, D, MS) and occupies a wide range of position along the anterior-posterior (A-P) axis of the animal. A UMAP of BWM cells identified distinct trajectories for head and non-head BWM (**Fig. 4E**). The non-head trajectory was formed by input trajectories that corresponded to lineages and progressed in parallel along the temporal axis. Using marker genes that are expressed in domains along the A-P axis (*15*, *33–35*), we divided BWM cells in the UMAP into six “bands” (**Fig. 4E**) and identified the specific anatomical cells present in each band (**Fig. 4F**, **Table S8**). We found that the Jensen-Shannon (JS) distance, a measure of transcriptome difference, between the transcriptomes of posterior (C lineage) vs. anterior (D/MS lineage) BWM cells did not decrease over time (**Fig. 4G**), indicating that BWM heterogeneity persists throughout differentiation.

## Temporal dynamics of the lineage-transcriptome relationship

We observed that the parent of a terminal neuron or glia was sometimes included at the beginning of the terminal cell trajectory, but the grandparent was almost never connected, with the exception of a few clonal lineages that produce only ciliated neurons (**Fig. 3A, S9, S10**). These discontinuities in trajectories led us to hypothesize that the final or penultimate cell division in a given lineage is frequently accompanied by a rapid transition from a transcriptome that is primarily shaped by lineage to one that is primarily shaped by cell fate.

We tested this hypothesis by measuring the similarity between transcriptomes of cells with different lineage relationships by using the JS distance. We focused our analysis on the AB lineage (ectoderm), which produces ∼70% of terminal cells in the embryo and undergoes roughly synchronized cell divisions. Most AB sub-lineages do not adopt a terminal fate until after the final round of division.

Using our lineage annotations (**Fig. 2**), we grouped AB lineage cells by their “cell generation,” i.e. the number of cell divisions since the AB founder cell. In AB generation 6 (“AB6”; embryo time ∼130-170 minutes), the transcriptomes of both sister and cousin cells tend to be similar (median JS distance = 0.28, **Fig. 5A**). In AB7 (∼170-230 min), lineage distance becomes significantly more predictive of transcriptome distance. Sister transcriptomes are more similar than cousins (p = 0.0035, Wilcoxon rank sum test), and sisters/cousins are more similar than cells from distal lineages (median JS distance = 0.26, 0.29, and 0.39 for sisters, cousins and 4th cousins respectively). Only one cell division later, in AB8 (∼230-300 min), the correlation between lineage distance and transcriptome distance weakens considerably. Another sharp transition occurs in AB9 (>300 min), which is the final round of division for most AB cells. Sister transcriptomes diverge sharply compared to the previous generation (median JS distance = 0.53 in AB9 vs. 0.31 in AB8), as do those of 1st-4th degree cousins. At this stage, lineage distance is almost completely uninformative for predicting transcriptome distance.

**Fig 5.**
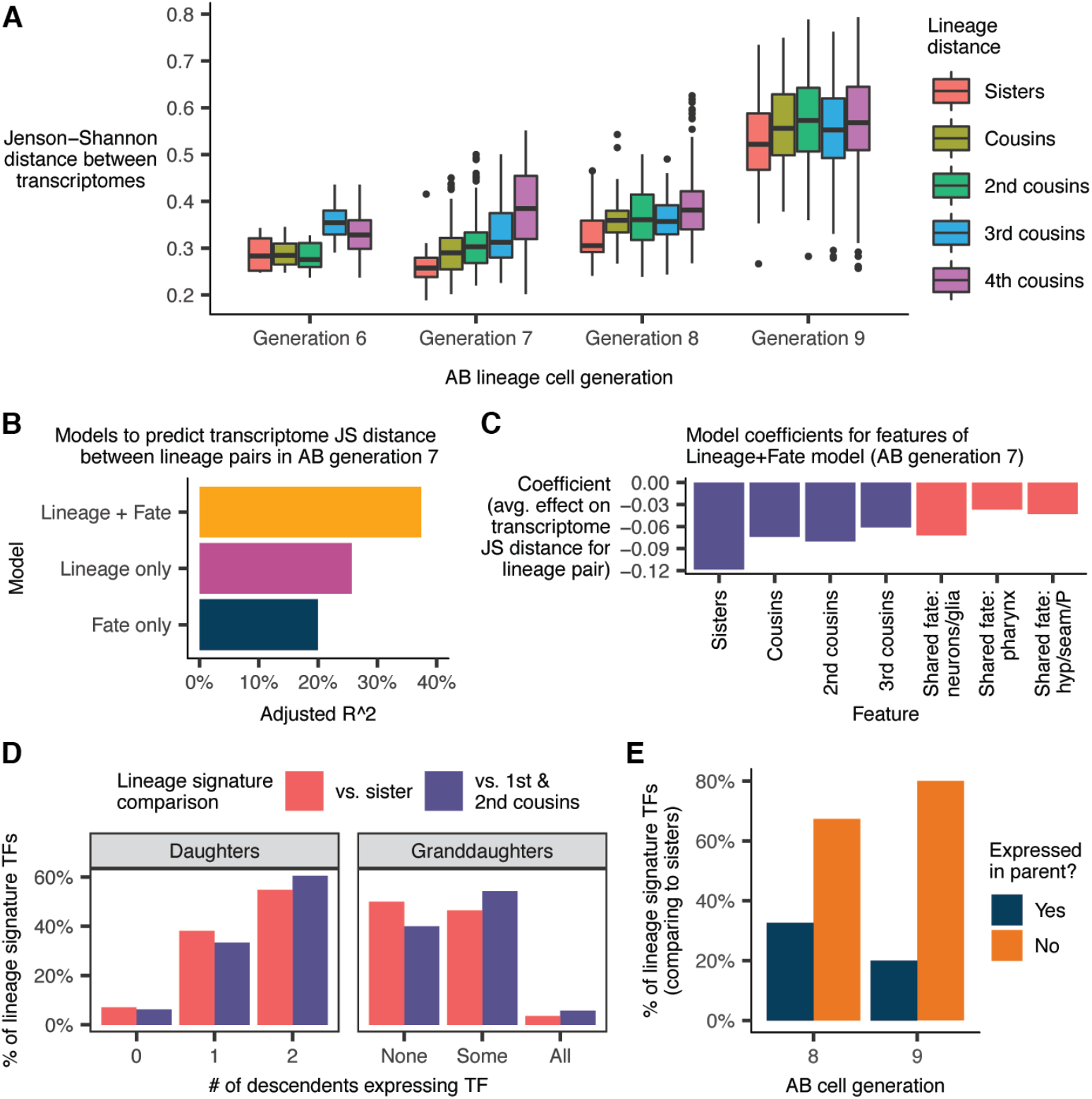
Correlations between cell lineage, cell fate, and the transcriptome. (**A**) Jensen-Shannon (JS) distance between the transcriptomes of pairs of AB lineage cells, faceted by cell generation and lineage distance. The “transcriptome” of a given anatomical cell is defined as the average gene expression profile of all sc-RNA-seq cells annotated as that anatomical cell (**Table S7**). (**B**) Performance of linear regression models to predict transcriptome JS distance between pairs of cells in AB generation 7. “Lineage” and “Fate” refer to sets of binary features (see text and **Methods**). (**C**) Model coefficients from the “Lineage + Fate” model. (**D**) Proportion of “lineage signature transcription factors”—TFs that distinguish cells in AB generation 7 from their sisters/cousins (see text and **Methods**)—that are expressed in the cells’ daughters and granddaughters. (**E**) Proportion of lineage signature TFs for cells in AB generations 8 and 9 that were previously expressed in the cell’s parent.

Most AB7 cells are multipotent, but their descendants usually share a tissue / broad cell type, e.g. epidermis vs. pharynx vs. neuron/glia. In the AB7 generation, lineage distance between cells was predictive of transcriptome distance (**Fig. 5A**). We sought to determine whether this trend was independent of, or redundant to, the trend of most AB7 cells being already committed to a tissue fate. A linear regression model to predict transcriptome distance between pairs of AB7 cells based on lineage distance and tissue fate explained significantly more variance than a model based on lineage or fate alone (**Fig. 5B-C**, also see **Methods**). This result suggests that the relationship between lineage and the transcriptome observed in **Fig. 5A** is not simply a by-product of lineage clades sharing a tissue fate.

Lineage does not predict the transcriptome well in AB8 or AB9, but this failure does not necessarily mean that genes characteristic of specific lineages in AB7 are no longer expressed. We considered the possibility that the transcriptomic “lineage signature” persists, but is simply overshadowed by differentiation-associated genes in terms of shaping the global transcriptome. To investigate, we first identified for each cell in AB7 a set of lineage signature transcription factors (TFs), which we defined as TFs that are both robustly expressed in the cell and enriched in that cell compared to either (a) its sister or (b) its first and second cousins (see **Methods**). There were on average 5.6 lineage signature TFs distinguishing a cell in AB7 from its sister, and 4.0 TFs distinguishing it from its 1st/2nd cousins. Next, we quantified how many of these lineage signature TFs were expressed (lower bound of 95% bootstrap confidence interval for expression level > 0) in the daughters or granddaughters of the AB7 cell with which the TF is associated.

The majority of lineage signature TFs for cells in AB7, compared to their sisters, were expressed in both of their daughters (**Fig. 5D**). A substantial minority (38%), however, were selectively inherited in just one daughter. Selective inheritance continues over the next division. 46% of lineage signature TFs for cells in AB7 were expressed in some, but not all of the cells’ annotated granddaughters (vs. 4% expressed in all granddaughters and 50% not expressed in any granddaughter). These statistics suggest that selective inheritance of TF expression patterns, previously observed in ciliated neuron lineages (**Fig. 3D, S15**), is a pervasive mode of gene regulation that contributes to the differentiation of sister lineages.

Sister lineages are also differentiated by the expression of new TFs characteristic of their terminal cell fate. 67% of lineage signature TFs that distinguish cells in AB8 from their sisters are not expressed in their parents; for AB9, this proportion increases to 80% (**Fig. 5E**).

## Global patterns of gene expression and transcriptome specialization

Hierarchical clustering of expression levels in all annotated lineages and cell types (**Tables S6 and S7**) provides a global view of expression dynamics for all genes in our dataset. A heatmap of pre-terminal lineage expression profiles (**Fig. S18**) does not reveal large clusters of genes specific to specific lineages, other than one cluster of genes specific to the early C and D lineages. Similarly, most marker genes used for lineage annotation are not part of large clusters of co-expressed genes. The clusters that do form are composed of early tissue-specific genes. The lack of cluster structure in the heatmap suggests that differential fates for tissue sub-lineages are specified by relatively small sets of genes. By contrast, a heatmap of terminal cell type expression profiles (**Fig. S19**) has more obvious structure. Cells in each major tissue express ∼500-1500 tissue-enriched genes. There is little reuse of tissue-enriched genes between tissues other than hypodermis, which shares many genes with glia and intestine. Neuron subtypes and other specialized cells (such as the hmc or M cell) are typically distinguished from other cells within their tissue by expression of <20-300 genes. Finally, there are substantial temporal changes in expression, especially in muscle and hypodermis.

We observed substantial variation between cells in the Gini coefficient, which measures how unequally different genes are expressed in a given cell type (**Fig. 6A**). Hypodermis, seam cells, and the pharyngeal gland express small sets of cell type specific genes at very high levels (high Gini coefficient), while the intestine and germline feature more diverse gene expression patterns (low Gini coefficient). In several cell types, increases in Gini coefficient over time coincide with decreases in the number of TFs expressed per cell (**Fig. S20A-C**). Families of TFs also exhibit differential expression patterns over time and across lineages. Nuclear hormone receptors (NHRs) are on average activated later in development than other TF families, such as Forkhead and Homeodomain TFs (**Fig. S20D**). Hypodermis and intestine express many distinct NHRs, while expression of Sox family TFs is largely restricted to neurons, glia and pharynx (**Fig. 6B**).

**Fig 6.**
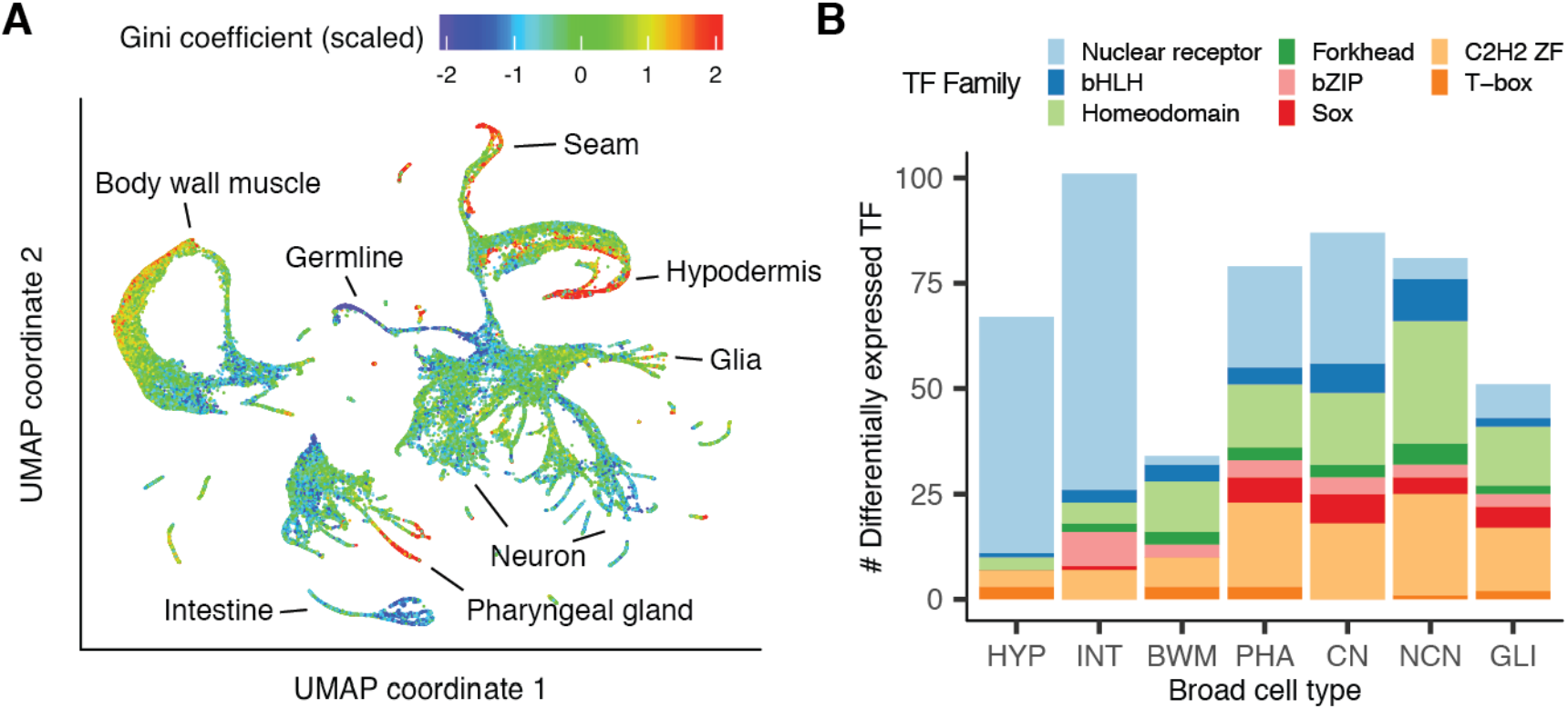
Transcriptome specialization and transcription factor usage across cell types. (**A**) A global UMAP with cells colored by the Gini coefficient of their gene expression vector, adjusted to correct for sample size bias and scaled by converting to z-scores. High Gini coefficients indicate that a small set of genes produces a large fraction of cell mRNA content. (**B**) Number of differentially expressed TFs and TF family composition across broad cell types. TF family annotations are taken from the CIS-BP database (41). Families that have fewer than 10 members detected in the current dataset were excluded from this plot.

## An RShiny app to explore and extend on our analysis

We developed VisCello, a Shiny app designed to distribute single cell analyses and provide interactive visualizations (**Fig. S21**). It is available as a web app (https://cello.shinyapps.io/celegans/) and can also be installed as an R package (https://github.com/qinzhu/VisCello). VisCello hosts dimensionality reductions (e.g. UMAPs), cell annotations, and marker gene tables for the different subsets of the data described in this manuscript. Users can visualize gene expression on UMAP or PCA plots, or as box/violin plots grouped by cell type or lineage. The plots are interactive, allowing users to zoom in on subsets of cells, define new cell annotation groups, and run differential expression analysis and GO/KEGG enrichment with these newly defined groups. Program state can be downloaded and shared, facilitating collaboration. VisCello can also be used to host and disseminate other single cell datasets (https://github.com/qinzhu/VisCello.base).

## Discussion

The cells of *C. elegans* are limited in number and invariant in lineage and cell fate, making it feasible to conduct comprehensive, whole-organism investigations. Yet within this limited repertoire of cells exists an impressive diversity of cell types, which work together to produce complex anatomical structures and behaviors. This study and our previous work (*2*, *15*) have shed light on the molecular basis for the specification of these cell types, but are only the first step toward a comprehensive understanding of the molecular basis of development. We hope that this resource will help guide future projects in the *C. elegans* community.

This dataset is unique compared to developmental sc-RNA-seq datasets from other species in that it links gene expression trajectories to the exact cell lineages they correspond to, allowing steps in the process of differentiation to be associated with specific cell division events. Thus, our data provide a quantitative portrait of Waddington’s landscape for a whole organism. The abruptness of many cell fate decisions in *C. elegans*, with many distinct terminal cell types becoming distinguished only in the final embryonic cell division, contrasts however with the smooth landscape in Waddington’s illustrations and warrants further investigation.

We observe convergence of gene expression patterns in many instances where distinct cell lineages produce identical or related cell types. Data from a recent atlas of mouse organogenesis (*11*) suggests that this phenomenon is also prevalent in vertebrates. For example, myocytes in the mouse atlas are produced by two convergent trajectories, and excitatory neurons are produced by several trajectories.

Our analysis suggests that it will be difficult to accurately connect developmental trajectories that start after the convergence of lineages with similar cell fates to trajectories that span earlier stages of development. A naive interpretation of the UMAP projection of the full dataset (**Fig. 1A**) could lead to inferred trajectories that are inconsistent with the correct lineage (for example, incorrectly concluding that hypodermis and seam cells are produced from a common ancestor that previously diverged from the progenitors of neurons). We linked progenitors to terminal cells using an iterative projection approach to increase resolution, and an extensive lineage-resolved imaging dataset from EPiC (*15*). No equivalent database exists for vertebrates, where detailed cell lineages are not known. Thus, we anticipate that to construct end-to-end trajectories of vertebrate organogenesis will require new lineage tracing experiments, such as using high-throughput CRISPR (*36*) or transposon (*9*) based methods. It will also require improved computational methods that can model heterogeneity among poorly-differentiated progenitor cells and highly-differentiated cell types in an integrated manner.

Between this study, our previous study of the L2 stage (*2*), and earlier studies of the 1 to 16-cell stage embryos (*37*, *38*), a large portion of the early *C. elegans* life-cycle has now been profiled by single cell transcriptomics. However, more datasets will be needed to complete missing stages, including other larval stages and the adult soma and germline. In the future, single cell profiling of different strains or species will be a useful approach to examine the evolution of cell types and their expression programs. All of these datasets will ideally be integrated into a single visualization platform, such as one based on VisCello, to allow full tracking of cell trajectories from fertilization through the end of life. A greater challenge will be to discover the precise mechanisms that produce transcriptomic outputs. Single cell transcriptome analysis of mutants will likely need to be integrated with new single cell multi-omic technologies (*39*) to bring mechanistic studies to a whole-organism scale.

## Acknowledgements

The raw data have been deposited with the Gene Expression Omnibus (www.ncbi.nlm.nih.gov/geo) under accession code GSE126954. This work was funded by NIH grants U41HG007355 and R01GM072675 to RHW, and R35GM127093 and R21HD085201 to JIM. This work was also funded in part by Commonwealth of Pennsylvania Health Research Formula Funds to JK, and by the William H. Gates Chair of Biomedical Sciences (RHW). We thank members of the Murray, Waterston and Kim labs, and Ben Lehner and Meera Sundaram for providing critical comments on the manuscript. We also thank Amanda Zacharias, Dionne Vafeados, Mitch Corson, Robert Terrell, Louis Gevirtzman and Peter Weisdepp for their contributions to the EPiC database. This article was prepared while HD was employed by the University of Pennsylvania. The opinions expressed in this article are the authors’ own and do not reflect the view of the National Institutes of Health, the Department of Health and Human Services, or the United States government.

## Materials & Methods

### Sample preparation

To obtain a broad range of embryo ages, including early stages, roughly synchronized C. elegans adults (N2 strain) were obtained by releasing embryos with standard hypochlorite treatment and letting the L1 larvae hatch and undergo growth arrest on unseeded plates. Starved L1s were transferred to NGM plates seeded with E. coli OP50 bacteria. Embryos were released from these synchronized young adults using hypochlorite treatment followed by three washes with L15-10 media. To generate cell suspensions, embryos were then treated with 0.5mg/ml chitinase at room temperature until the shells were dissolved (30-40 minutes at room temperature (∼22 degrees C)) followed by dissociation of the cells using a 3ml syringe fitted with 21 1/2 gauge needle until >80% of embryos were disrupted. The cell suspension was then passed through a 10μM filter, washed in phosphate buffered saline (PBS) and finally resuspended in PBS. An estimated 14,000 cells were loaded immediately on a 10X Chromium instrument. The trypan blue negative viable cell count was estimated using a hemocytometer and was >84% for all samples.

To sample later stages more deeply, more tightly synchronized embryo populations (used for the 300-minute, 400-minute, and 500-minute time series shown in **Fig. 1B**) were obtained through two cycles of bleaching adult worms (strain VC2010, a strain derived from N2 that has been completely sequenced). On the first round of synchronization, populations of mixed stage embryos recovered by hypochlorite treatment of mixed populations were hatched overnight in egg buffer (118 mM NaCl, 48 mM KCl, 3 mM CaCl2, 3 mM MgCl2, 5 mM HEPES pH 7.2) with gentle shaking. The hatched L1s were plated onto 150 mm peptone rich NGM plates seeded with E. coli NA22 at no more than 100,000 worms per plate. When worms reached the adult stage, the number of embryos inside the adults was monitored until most had about 4 embryos on each gonad arm. The adult worms were collected and treated with hypochlorite to release embryos. The embryos were again allowed to hatch in the absence of food at 20 °C for 12 hours yielding a more tightly synchronized population of L1 worms. Around 250,000 L1 larvae were plated onto four 100 mm petri plates seeded with NA22 bacteria and allowed to develop at 20 °C. As the worms reached the young adult stage, the population was closely monitored. When about 20-30% of the adults had a single in either arm of the gonad, worms were subjected to hypochlorite treatment. The time hypochlorite was added to the worms was considered t=0 (see Warner et al. in press, for typical age distributions). The capture time was taken as when the cells were loaded onto the 10x Chromium capture. The embryos were allowed to develop in egg buffer until one hour prior to capture time. The embryos were collected by centrifugation, resuspended in 0.5 ml egg buffer and 1 ml chitinase (1 U/ml) and transferred to 30 mm petri dishes. The degradation of eggshell was monitored; after ∼20 min (about half the eggs had lost the shell), the suspension was transferred to a 15ml falcon tube and centrifuged at 200 g for 5 min. The chitinase solution was aspirated; a solution of 200 ul pronase (15mg/ml) together with 0.5 ml egg buffer was added to the embryo pellet. The vitelline membrane was disrupted and the cells released by repeated passage through 21G1 ¼ needle attached to a 1 ml syringe. When sufficient single cells were observed the reaction was stopped by adding 1ml of egg buffer containing 1% BSA. Cells were separated from intact embryos by centrifuging the pronase treated embryos at 150 g for 5 min at 4°C. The supernatant was transferred to a 1.5 ml microcentrifuge tube and centrifuged at 500 g for 5 min at 4°C. The cell pellet was washed twice with egg-buffer containing 1% BSA.

Single cell capture and library preparation followed 10X Genomics published protocols. For each channel, 14,000 C elegans cells were mixed with reverse transcriptase reaction solution and loaded immediately onto the capture chip to minimize the time C. elegans cells spent in reverse transcription cocktail. The exception was the first 500 minute sample, when 14,000, 4,666, and 1,555 cells were loaded on each channel.

### Read mapping and gene expression quantification

The single cell RNA-seq data was processed using the 10X Genomics CellRanger pipeline. Reads were mapped to the C. elegans reference transcriptome from Wormbase, version WS260. We noticed that many 3’ UTR annotations in the reference transcriptome were too short, causing genic reads to be called as intergenic, affecting gene expression quantification. To address this, we also mapped reads to modified versions of the WS260 transcriptome in which all 3’ UTRs were extended by either 100, 200, 300, 400, or 500 bp (these 3’ UTR extensions were cut short if the extended UTR would overlap with a downstream gene).

We then defined a set of criteria that specified for each gene whether it was beneficial to extend the 3’ UTR for that gene, and if so, by how much. For each gene, we counted the number of reads across the entire dataset mapped to that gene for each version of the reference. We computed the ratio of the read counts from the 500 bp 3’ UTR extended reference to the baseline reference. If this ratio was < 1.2, or if the total read count for the gene in the 500 bp 3’ UTR extended reference was < 20, we used the baseline 3’ UTR annotation for that gene. Otherwise, we used the shortest 3’ UTR extension (100, 200, 300, 400, or 500 bp) that gave at least 90% of the read count gain that was given by the 500 bp 3’ UTR extension.

We repeated this process with reads from our previous study on L2 worms (*1*). If a gene met our criteria for extending the 3’ UTR based on embryo reads, we used the extension length determined by the embryo reads. If a gene did not meet our criteria for extending the 3’ UTR based on embryo reads but did meet the criteria based on L2 stage reads, we used the extension length determined by the L2 stage reads. After deciding on how much to extend each gene’s 3’ UTR, we made a final reference transcriptome incorporating all of the per-gene 3’ UTR extension lengths. We then used this final reference transcriptome as input to the CellRanger pipeline to generate gene-by-cell UMI count matrices.

Our final reference transcriptome is available as **Supplemental File 1**. It should be suitable for future studies of *C. elegans* embryos or early larva. However, neither the embryonic or L2 datasets contain reads from developed germline cells (sperm and oocytes) or developed somatic gonad cells (e.g. spermatheca), so our reference will not properly extend 3’ UTRs for genes specific to these cell types.

### Criteria for distinguishing cells from empty droplets

The default barcode filtering algorithm in the 10X CellRanger pipeline can fail for experiments where the cells profiled are highly variable in size, resulting in a non-normal distribution of UMIs per cell. This is the case for our data. The total volume of the *C. elegans* embryo remains constant as cells divide within it, making cells of later generations smaller than those from earlier generations. Additionally, some cell types are more prone to damage and mRNA leakage than others. Neurons in particular usually have lower UMI counts than other cell types. To account for these factors, we manually set UMI count thresholds to distinguish cell barcodes from empty droplet barcodes on a sample-by-sample basis, based on the knee plots reported by CellRanger. The UMI count thresholds ranged for 700-1100.

While performing downstream analyses, we noticed that several neuron types were missing from our data. We discovered this was due to neuron cells with extra low UMI counts (< 700 UMIs) being excluded by our UMI count thresholds. Lowering the UMI count threshold for all cells however would include low-quality, potentially damaged cells for other cell types where the average UMIs/cell is larger. To integrate the low-UMI count neurons, we:

1. made a set of all cells with UMI count >= 500 (vs. the previous threshold of 700)
2. ran UMAP dimensionality reduction (described below) on this set of cells
3. identified clusters of cells corresponding to neurons using the marker genes *sbt-1* and *egl-21*
4. ran UMAP again for the just the neuron cells
5. filtered putative doublets (i.e. cells also expressing markers of non-neuronal cell types)
6. made whitelists of the remaining cells

These whitelisted low-UMI count neurons were then included when generating the final neuron UMAPs presented in this paper (**Fig. 3A, S10**).

### Dimensionality reduction

For each dimensionality reduction (both for the global analysis of all cells and the tissue specific analyses), the first step was to perform PCA and adjust the PCA results to correct for batch effects. We performed PCA on the size-factor corrected, log transformed expression matrix, typically with 50-100 PCs depending on the dataset.

For batch effect correction, we noted that the predominant source of batch effects in our data appeared to be background contamination where RNA from lysed or damaged cells enters droplets in the 10X sc-RNA-seq apparatus that contain intact cells, causing each cell to receive reads from exogenous RNA. For each experimental sample, we computed the gene expression distribution of this background RNA by summing the read counts for cell barcodes that had < 50 UMI, i.e. empty droplets. We transformed the background RNA count vector for each sample as if it were the count vector for a cell, and projected this vector into the PCA space computed from real cells. We then computed the dot product of each real cell PCA coordinate vector with each sample’ background vector, calling this the “background loading” of a given cell for a given sample (each cell actually comes from exactly one sample, but computing each cell’s loading for each samples’ background made the next step mathematically/computationally simpler). Next, we fit a linear regression model—real cell PCA coordinate matrix ∼ cell background loadings—and called the residuals of that model the “background corrected PCA matrix.” This background correction method is similar to, but developed independently of, a recently published method (*2*).

We reduced the dimensionality of the background corrected PCA matrix to 2 or 3 dimensions using the UMAP algorithm (*3*, *4*), using the wrapper function for this algorithm provided by the Monocle software package, version 3 alpha (the reduceDimension function). The UMAP parameters were: metric = “cosine”, min_dist = 0.1, n_neighbors = 20. Lastly, cells in the UMAP space were clustered using the Louvain algorithm (*5*).

### Doublet identification

We used two complementary methods to identify doublets. The first method involved identifying clusters of doublets in iterated UMAP projections of the data on the basis of coexpression of high-confidence cell type specific marker genes, reported in Wormbase (*6*), for >1 cell type (e.g. a cluster expressing the muscle markers *myo-3* and *pat-10* along with the neuron markers *egl-21* and *sbt-1* was considered a muscle-neuron doublet cluster). We applied this simple approach to a global UMAP of all cells and iterated UMAPs of tissues / related groups of cells from the global UMAP (i.e. muscle, intestine, ciliated neurons, etc.).

The second approach involved logistic regression models, one for each broad terminal cell type class (i.e. muscle, intestine, ciliated neurons, etc.), that predict whether a cell is part of that cell type class or not. We fit one such model for each broad cell type class and used the models to score each cell for the probability of it being a member of each broad cell type class. Cells that had >= 2 cell types with a >= 20% predicted probability of the cell being a member of that cell type were considered doublets. Clusters in the UMAP projections that were enriched for cells considered doublets by these regression models were manually examined, and in some cases manually filtered.

Due to the abundance of cell type specific marker genes, we estimate that we were able to filter out almost all terminal cell type doublets. Residual expression of genes from one cell type in a cluster corresponding to another cell type appears to be driven by background RNA contamination, not doublets. Our approach is less likely to catch doublets between early stage cells that do not yet express marker genes of differentiated terminal cell types. For early stage embryos however, the cell dissociation protocol works more reliably than for late stage embryos, so we expect the doublet rate to be close to the reported rate for the 10X Genomics Chromium platform, which is low (∼4.5% given ∼9k cells loaded per lane).

### Embryo time estimation

For each cell, we estimated the age of the embryo that the cell came from (“embryo time”) based on Pearson correlation of its transcriptome with a bulk RNA-seq time series data from Hashimshony *et al*. (*7*). Their data show that the majority of temporal genes expressed in any given lineage are not lineage specific. Thus, we first defined a list of genes with time-dependent expression patterns, requiring an auto-correlation greater than 0.6 and standard deviation greater than 1.5 across bulk time points (units = log TPM). Pearson correlation was then computed between log-scaled single cell and bulk data using only the time-dependent genes. We observed for non-multiplet cells, the Pearson correlation across time show strong peak pattern (**Fig. S1A**). Thus, by fitting a loess regression curve and finding its maximal point, we were able to assign each cell with its most correlated bulk time point.

The time estimated based on data from Hashimshony *et al* approximately agrees with the collection time (**Fig. S1B**), and has strong correlation with that estimated based on data from Boeck *et al* (**Fig. S1C**). Furthermore, as the live imaging data provide an accurate cell birth time estimation, we correlated 5th percentile of estimated embryo time within each early lineage cell bin to the corresponding birth time, and found significant correlation between the two (**Fig. S1D**). However, we also noticed that the estimated time does not exactly match the birth time, and is skewed towards earlier time points. This is likely due to variation between different platform and techniques, as single cell data contains more drop-outs compare to bulk RNA-Seq. We compared our estimated embryo time to those inferred by some of current pseudotime analysis methods (data not shown), and found only the estimated embryo time consistently matches the complex differentiation trajectories in the UMAP.

In the Waterston lab samples, embryos were incubated for a specific amount of time after hypochlorite treatment. However, each sample has some outlier cells with abnormally low embryo time estimates, i.e. lower than the incubation time. There are several biological and technical factors that could produce these outlier cells. The developmental rate of *C. elegans* embryos can vary by over 2-fold depending on temperature, and may also be influenced by differences in crowding, hypoxia, or the effects of hypochlorite and chitinase treatment. Consistent with this, embryo time measurements estimated using the Boeck et al dataset, which was collected using methods more similar to those used in this study, were systematically later than embryo time estimated from the Hashimshony data (**Fig. S1C**). Alternatively, some cells may have embryo time estimates that are lower than the true developmental age of the embryo they came from. Sparsity in the single cell data contributes to noise in the estimates. Finally, the most extreme outlier embryo time estimates in each sample are for germline cells. The germline maintains expression of many genes that turn off during early embryogenesis in all other cells. This causes embryo time estimates based on correlation to bulk RNA-seq to be inaccurate for this cell type.

### Per-cell background correction and filtering

Our method for correcting for background RNA contamination, described in the “Dimensionality Reduction” section of this supplement, works solely on the level of PCA coordinates and does not change the underlying gene-by-cell expression matrices. We used a separate background correction method to adjust these gene expression matrices on a per-cell basis for purposes of making plots of gene expression.

Our per-cell background correction method relies on a panel of cell-type specific marker genes, that are assumed based on the literature (and confirmed empirically in our data) to be specific to either hypodermis (including seam and P cells) or body wall muscle (BWM). The hypodermis-specific genes were: *sqt-3, dpy-17, dpy-14, dpy-10, dpy-7, dpy-2, dpy-3, bus-8, wrt-2*, and *noah-1*. The BWM-specific genes were: *pat-10, mlc-3, cpn-3, clik-1, ost-1, mlc-1, mlc-2, tni-1, ttn-1, unc-15*, and *myo-3*.

The gene expression distribution for the background contamination of each biological sample was estimated by aggregating the reads for cell barcodes that had < 50 UMIs, which were assumed to correspond to empty droplets in the 10X sc-RNA-seq apparatus. The expression level of each gene in the panel was computed for each sample’s background, measured in transcripts per million (TPM). Similarly, the expression level of each gene in the panel was computed for each cell, also measured in TPM. The background fraction of a cell was estimated as the sum of the expression of panel genes in the cell divided by the sum of the expression of panel genes in the background distribution for the sample that cell came from. For cells annotated as hypodermis, glia, or potential progenitors of those cell types, hypodermis-specific genes from the panel were excluded from the computation. Likewise, for cells annotated as body wall muscle, intestinal/rectal muscle, or a non-pharyngeal mesoderm cell type, as well as progenitors of those cell types, BWM-specific genes from the panel were excluded from the computation. For all other cells, all genes from the panel were used.

The median estimated background fraction across all cells in the dataset was 17.7%. Putatively damaged cells with an estimated background fraction >= 75% (8.3% of all cells, see **Fig. S22A**) were filtered entirely from all subsequent plots and analyses. For the remaining cells, the cell expression profiles were corrected to subtract the contribution from background. A cell’s raw gene expression vector (UMI counts) was converted to transcripts per million by dividing each entry by the sum and multiplying by one million. The background-corrected TPM value for each gene was computed according to the formula:

> background-corrected TPM = max(raw TPM – background fraction * background TPM, 0)

where background TPM is the expression of the given gene in the background distribution for the biological sample the cell came from. The background-corrected corrected TPM values were then rescaled to once again sum to 1,000,000 and then converted back into (pseudo-)counts based on the total UMI count of the cell. Fractional count values were rounded probabilistically (i.e. a value of 2.7 was rounded to 3.0 with a 70% chance and to 2.0 with a 30% chance).

After background correction, cells with low background fractions and cells with high background fractions have near-identical average gene expression profiles (**Fig. S22B**). This indicates that non-background gene expression observed in high background cells is not systematically biased compared to low background cells.

### Computing aggregate gene expression profiles for cell types and lineages

To compute the aggregate gene expression profile for a cell type (**Table S6**) or lineage (**Table S7**), we (1) subsetted the whole-dataset gene-by-cell gene expression matrix to include just the cells annotated as the given cell type or lineage; (2) divided each column by the corresponding cell’s size factor (a statistic computed by the Monocle software package equal to the cell’s total UMI count divided by the geometric mean of all cell’s total UMI counts); (3) took the mean of each row (gene); and (4) rescaled the resulting vector to sum to 1,000,000. This results in a gene expression vector measured in transcripts per million (TPM).

We performed these computations using the original gene expression matrix, not corrected for background RNA contamination. After computing the aggregate gene expression vector for a cell type or lineage, we then corrected it for background contamination using the same method described in the previous section (“Per-cell background correction and filtering”), treating the aggregate vector as if it were the expression vector for a single cell. Compared to the alternative option of correcting each cell for background first and then computing the aggregate profile, aggregating first then correcting makes the estimate of background fraction more robust due to an increased sample size (number of reads).

Even after background correction, we noticed some residual, aberrant expression of genes that should not be expressed in a given cell type. In several cases, this aberrant expression was due to just one or two outlier cells within a given UMAP cluster. We suspected these outlier cells included both doublets missed by our filtering procedure and “pseudo-doublets”, consisting of a real cell plus debris from another cell. In order to reduce the impact of these outliers, in **Table S6** and **Table S7** we report a “robust” estimate of the mean expression for a gene in a given cell type or lineage (in addition to the “raw” estimate). This robust estimate excludes the highest-expressing cell and the lower-expressing cell for a given gene in a given cell type / lineage before computing the mean expression.

The impact of outliers is greater for cell types represented by only a small number of cells in our dataset, and for cell types that have a low average number of UMI per cell. Estimates of mean gene expression values are therefore less precise for these cell types. To estimate the variance of our mean gene expression statistics, we used bootstrap resampling: for a cell type with N cells in our dataset, we randomly sampled, with replacement, N cells from that set and computed mean expression statistics from that sample. We repeated this process for 1,0 iterations for each cell type and computed bootstrap confidence intervals from the resulting distribution of mean estimates. If one must make a statement that a gene is “expressed” in a given cell type or lineage, we recommend using the criterion that the lower bound of the 95% bootstrap confidence interval is >0 TPM.

### Differential expression analysis for Fig. 3D and Fig. S15

We included four classes of transcription factors (TFs) in the heatmaps of **Fig. 3D** and **Fig. S15**. Both figures consider differential expression of TFs between different ciliated neuron lineages. For the division of a parent neuroblast into two daughter cells, the four TF classes of interest were:

1. TFs enriched in one daughter vs. the parent and vs. the other daughter
2. TFs depleted in one daughter vs. the parent and vs. the other daughter
3. TFs enriched in the parent vs. both daughters and vs. other neuroblasts of the same generation
4. TFs enriched in parent vs. other neuroblasts of the same generation; and in both daughters vs. other terminal cells

We considered a TF “enriched” in cell set A vs. cell set B if the expression in A vs at least 3-fold higher than in B; and if the difference in expression was statistically significant with q-value < 0.01. We considered a TF “depleted” in cell set A vs. cell set B if it was “enriched” in B vs. A. q-values were computed using Monocle’s differentialGeneTest function. Differential expression tests were performed for all genes, not just TFs—the non-TF results were discarded, but this was done to produce more conservative q-values compared to considering only TF DE tests. Cells with embryo time >650 minutes were excluded from all comparisons. Due to limited figure space, some TFs that matched the criteria of the four TF classes but had low absolute expression levels were excluded from the figure heatmaps.

### Derivation of lineage and fate signature genes for Fig. 4D

Lineage signature genes were derived by one vs rest differential expression analysis on the three input lineages based on Louvain clustering results and annotations from **Fig. S12**, using “sSeq” implemented in the cellrangerRkit package (*8*). Genes associated with IL1/IL2 terminal fate were derived by comparing IL1/IL2 cells to all other ciliated neurons in **Fig. 3A**. For each of the gene sets, the average TPM across all genes in the set was computed for cells from each of the three input branches, binned in 30-minute intervals up to 390 min, where the branches can no longer be distinguished from each other in the UMAP. Values in each heatmap were linearly rescaled to be within the range of 0 to 1.

### Regression models for Fig. 5B

In **Fig. 5B**, we show the performance (R^2) of a set of linear regression models. In each model, the data points are pairs of cells from generation 7 of the AB lineage (anatomical cells, not sc-RNA-seq cells) that are 4th cousins or closer in the lineage. The response of each model is the Jensen-Shannon distance between the average gene expression profiles of the two cells, as reported in **Table S7**. These models use one or both of two sets of binary features. The first set of features (“Lineage”) are binary flags indicating whether the cells being compared are sisters, cousins, 2nd cousins, or 3rd cousins. Pairs of cells that are 4th cousins will have none of these flag variables set to true. The second set of features (“Fate”) are binary flags for whether the lineages produced by each AB generation 7 cell are both (a) progenitors of epidermal cells (hypodermis, seam, and P cells) only, (b) progenitors of pharyngeal cells only, or (c) progenitors of neurons and glia only. Pairs of cells with mismatched fate, e.g. an epidermis progenitor and a pharynx progenitor, will have none of these flag variables set to true.

### Definition of lineage signature transcription factors for Fig. 5D and 5E

For the analyses presented in **Fig. 5D-E**, we introduce a concept of “lineage signature transcription factors.” For each anatomical cell that we annotate in the AB lineage, we define the set of lineage signature transcription factors associated with that cell to be set of TFs that satisfy both of the following criteria:

1. The lower bound of the 95% bootstrap confidence interval for the average expression level of the TF in the cell, as reported in **Table S7**, is >0. In other words, the TF must be robustly expressed in the cell.
2. The TF must be expressed at least 5-fold higher in the cell compared to its sister in the lineage, with a differential expression q-value < 0.01.

We also define an alternate set of lineage signature transcription factors for each cell, in which instead of comparing against the sister of a given cell in criterion (2), we compare against the average expression profile of all annotated 1st and 2nd cousins of the cell.

### Computing adjusted Gini coefficient for Fig. 6A

Gini coefficient is biased by sample size (*9*). Therefore, to adjust for total UMI count difference between cells, we first downsample counts of each cell to a total of 500 UMIs (the minimal UMI count across all cells) using a multinomial distribution, with probability equal to each gene’s UMI count divided by total UMI count of the cell. We then computed Gini coefficient on this adjusted vector, and used the z-score of the adjust Gini coefficient for comparing transcriptome inequality across cells.

## Supplemental Note 1

In the analysis of single cell RNA-seq data, the term “trajectory” is often used to refer to a group of cells that 1) represent a specific cell lineage or cell type, and 2) can be ordered in a way that reflects the cells’ progression over time from one transcriptomic state to another. Algorithms for trajectory inference can construct such orderings by mapping high dimensional gene expression data to a low dimension and fitting a graph structure to the data points in the low dimension. The distance between a user-defined root vertex on the graph to the location of a cell on the graph is called the cell’s “pseudotime,” and for a particular path along the graph, the ordering of cells on that path by pseudotime defines a “trajectory” (the graph as a whole can be considered a “branched trajectory”).

In this manuscript, we have not used any trajectory inference algorithm. Instead, we use our embryo time estimates for each cell, which are computed based on correlation of the cell transcriptome with a high-resolution bulk RNA-seq time series (see **Methods** and **Fig. S1**), as a universal ordering for all cells in the dataset. We annotate cell types and lineages based on marker genes from the literature (**Table S1, S4**) and on clustering in UMAPs of our data. Ordering cells with a common cell type or lineage annotation by embryo time defines a “trajectory”. In most cases, trajectories defined in this way also form contiguous shapes in UMAPs, to which one could fit a graph structure to if one wanted to. Our universal ordering of cells by embryo time is a more robust approach however, as 1) it allows annotations from multiple UMAPs to be integrated to define a single “trajectory” per cell type/lineage, and 2) the approach still works even when cells of a common type/lineage are split into disparate, noncontiguous groups in a UMAP (which may occur due to abrupt changes in gene expression or due to technical factors, such as non-uniform sampling with respect to time).

**Fig. S1.**
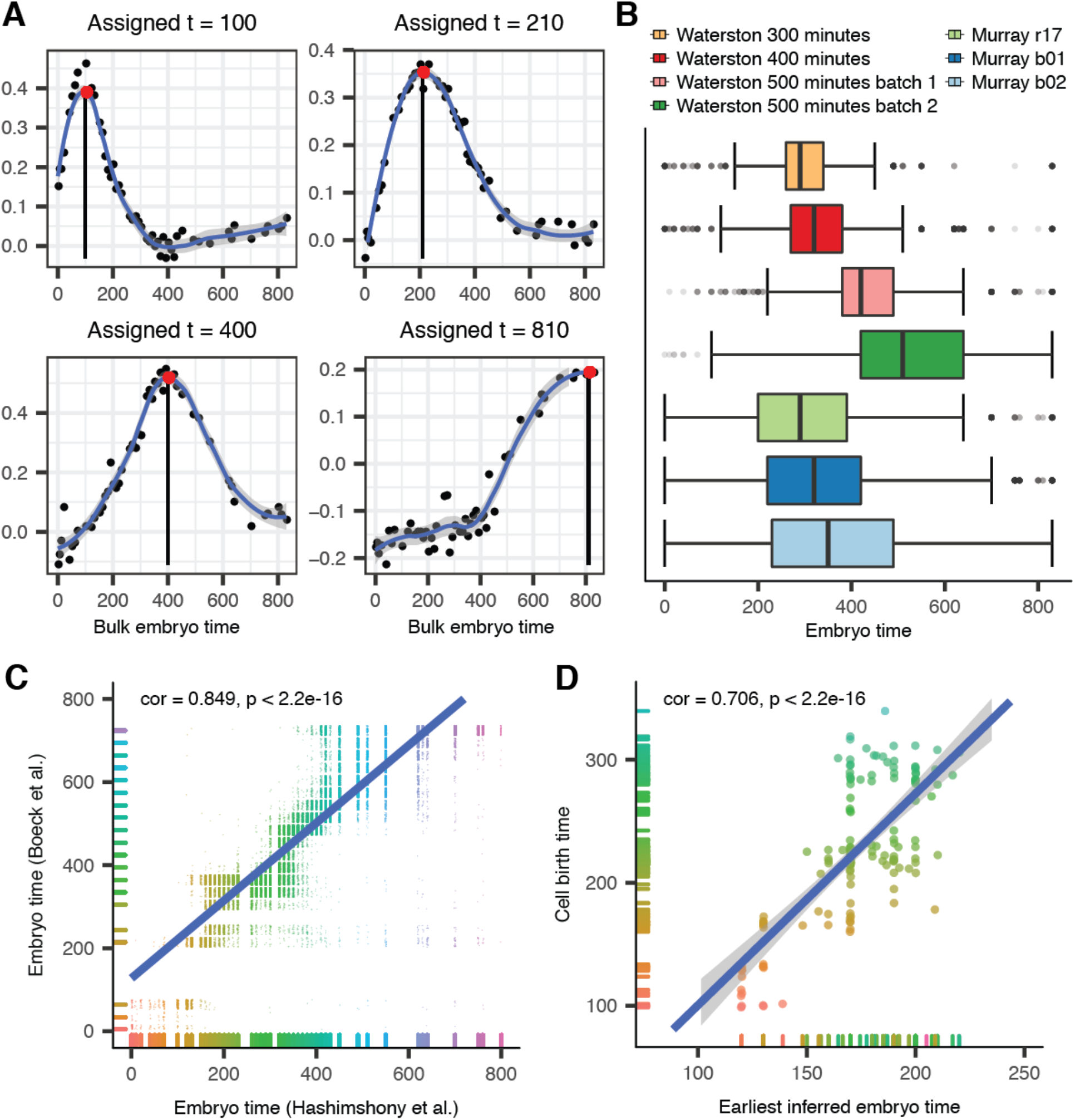
Method for estimating the age of the embryo an sc-RNA-seq cell came from. Embryo times are measured in minutes post first cleavage. (**A**) Embryo times are estimated based on Spearman correlation of a single cell’s transcriptome to a bulk RNA-seq time series. Pointwise estimates of the correlation to each time point are smoothed using a loess regression. (**B**) Distribution of estimated embryo times for each biological sample. The average embryo time estimate in the Waterston lab sample correlates with the real time duration that the embryos were incubated. Each sample contains some outlier cells with abnormally low embryo times. Potential biological and technical causes for the presence of these outlier cells are discussed in the **Methods**. (**C**) Correlation of embryo time estimates based on Hashimshony *et al*. (*7*) to an alternate set of embryo time estimates based on Boeck *et al*. (*10*). Estimates based on Hashimshony *et al* were used for all downstream analyses. (**D**) Correlation between cell birth times estimated based on our lineage annotations (x-axis) with cell birth times computed based on automated analysis of imaging data (y-axis) (*11*).

**Fig. S2.**
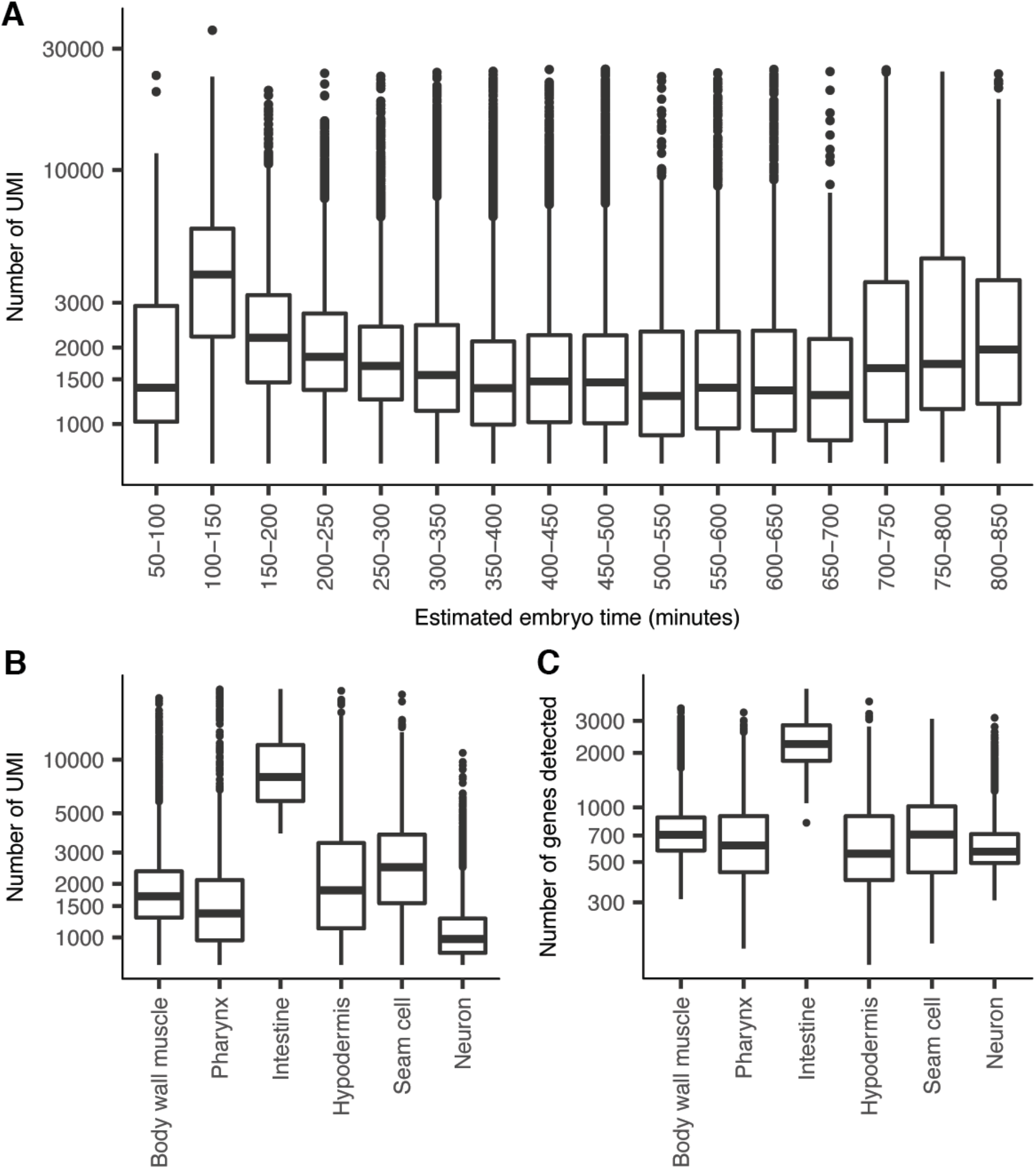
UMIs recovered per cell decreases with embryo age. All Y-axes are log scaled. (**A**) Distributions of number of UMIs recovered per cell, binned by estimated embryo age. Note that our quality control procedures exclude cells with < 700 UMIs (or < 500 UMIs for neurons), causing the decrease in UMIs/cell to be understated, as the proportion of cells falling below the cutoff is greater for later stage embryos. The apparent increase in UMIs per cell at >650 minutes is due to the cell type recovery bias in these late, cuticle-synthesizing embryos. (**B** and **C**) Number of UMIs and genes detected for cells with embryo time in [390, 650], by tissue.

**Fig. S3.**
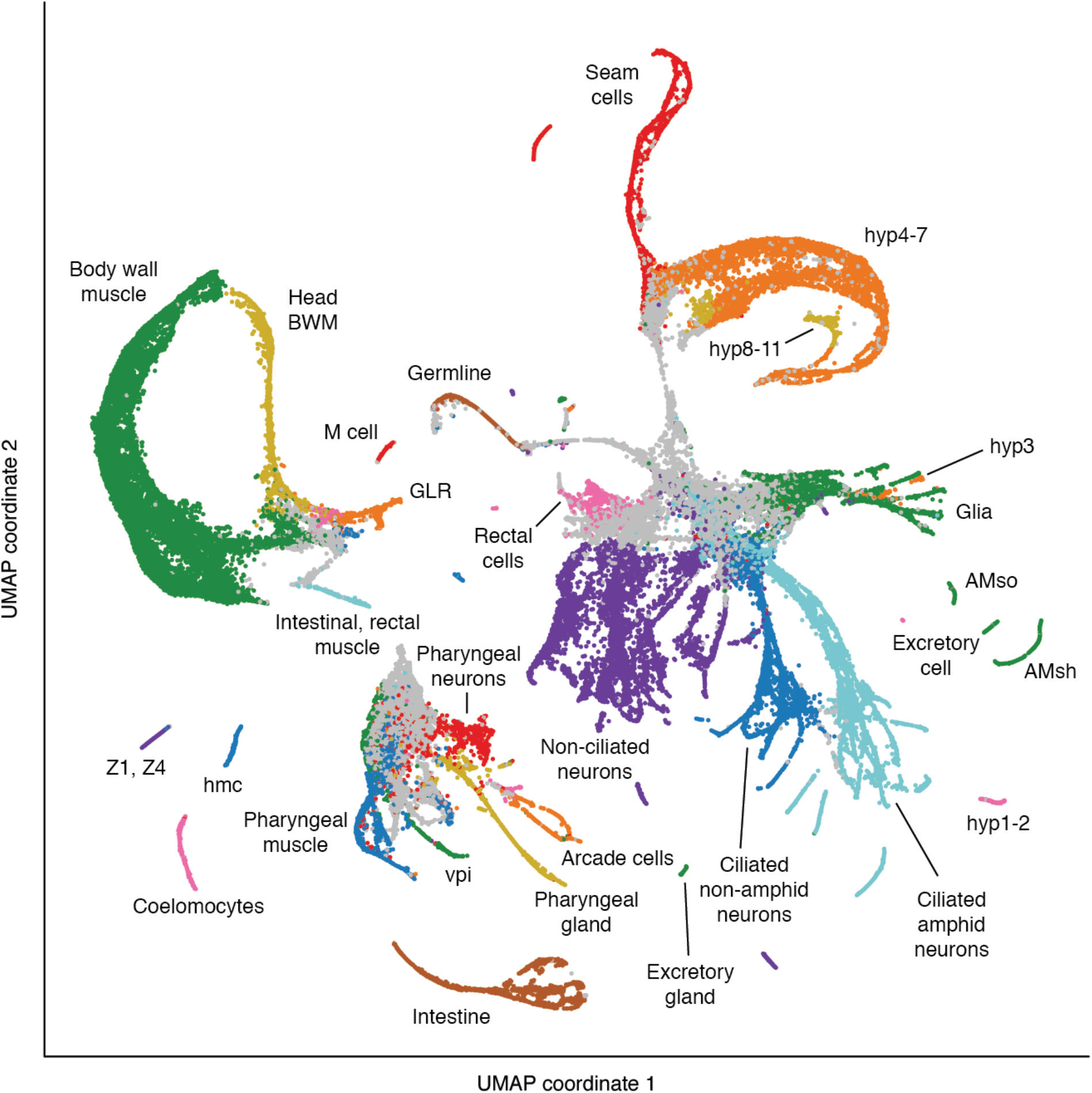
Cell type annotations for the global UMAP. This plot shows more cell type annotations for the global UMAP from **Fig. 1**. For fine-grained annotations of cell types in each major tissue, see **Fig. 3A** and **Fig. S5–11**. For fine-grained annotations of early cell lineages, see **Fig. S12–13**.

**Fig. S4.**
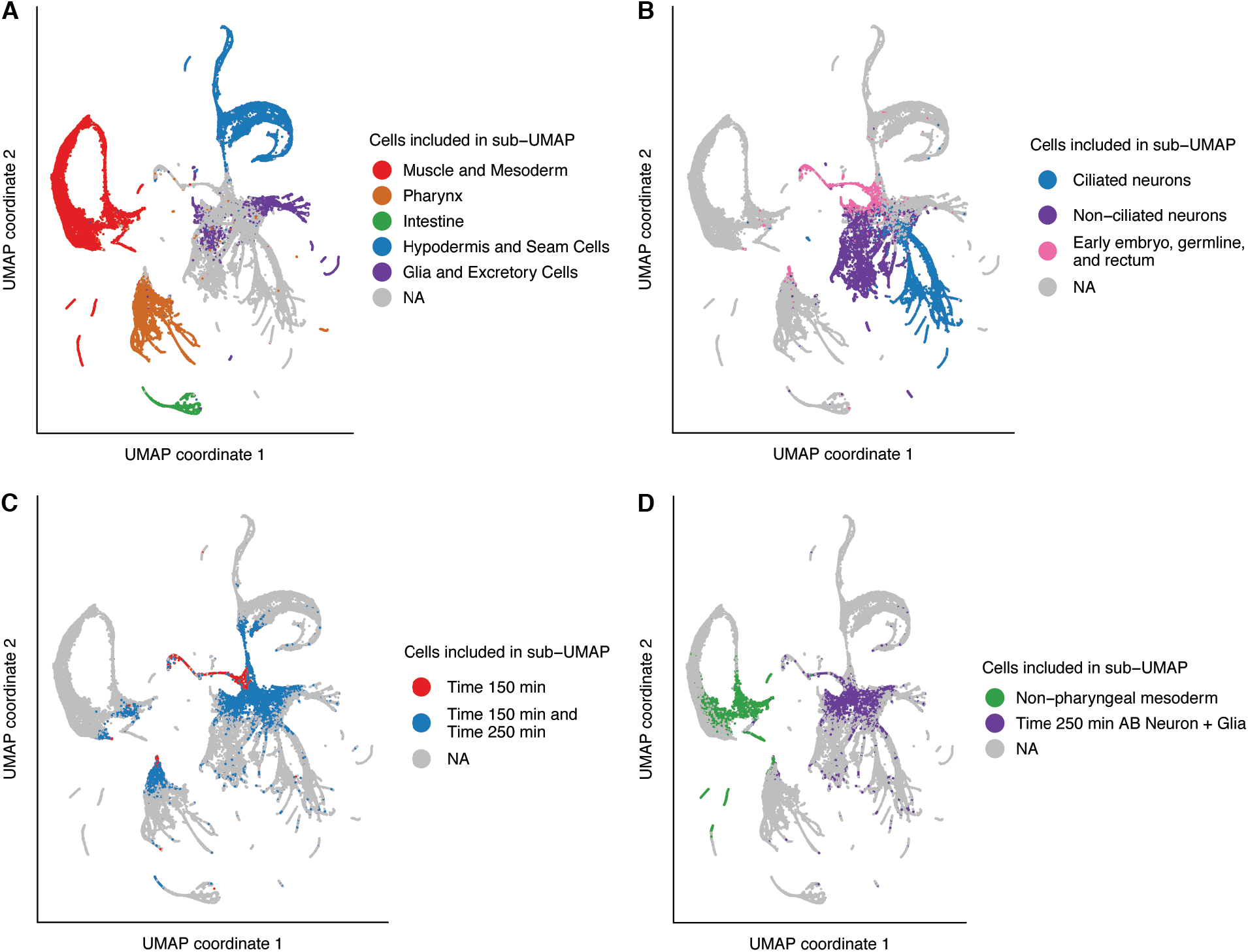
Cells included in each sub-UMAP. Plots show which cells from the global UMAP (**Fig. S3**) are included in each sub-UMAP, including UMAPs aimed at visualizing terminal cell types (**A, B**) and UMAPs focused aimed at visualizing early lineages (**C, D**). Plots of annotations in each sub-UMAP are shown in **Fig. S5–S12**. Note that the actual assignment of cells to sub-UMAPs was performed based on a 3D version of the global UMAP (not shown). In (**C**), all cells included in the Time 150 min. sub-UMAP are also included in the Time 250 min. sub-UMAP.

Note: The figures below show UMAPs of muscle and the non-pharyngeal mesoderm (**Fig. S5**), pharynx (**Fig. S6**), intestine (**Fig. S7**), hypodermis and seam cells (**Fig. S8**), glia and excretory cells (**Fig. S9**), non-ciliated neurons (**Fig. S10**), and germline and rectum (**Fig. S11**). A UMAP of ciliated neurons is shown in the main text (**Fig. 3A**). UMAPs of early stage cells are shown in **Fig. S12** and **Fig. S13**.

**Fig. S5.**
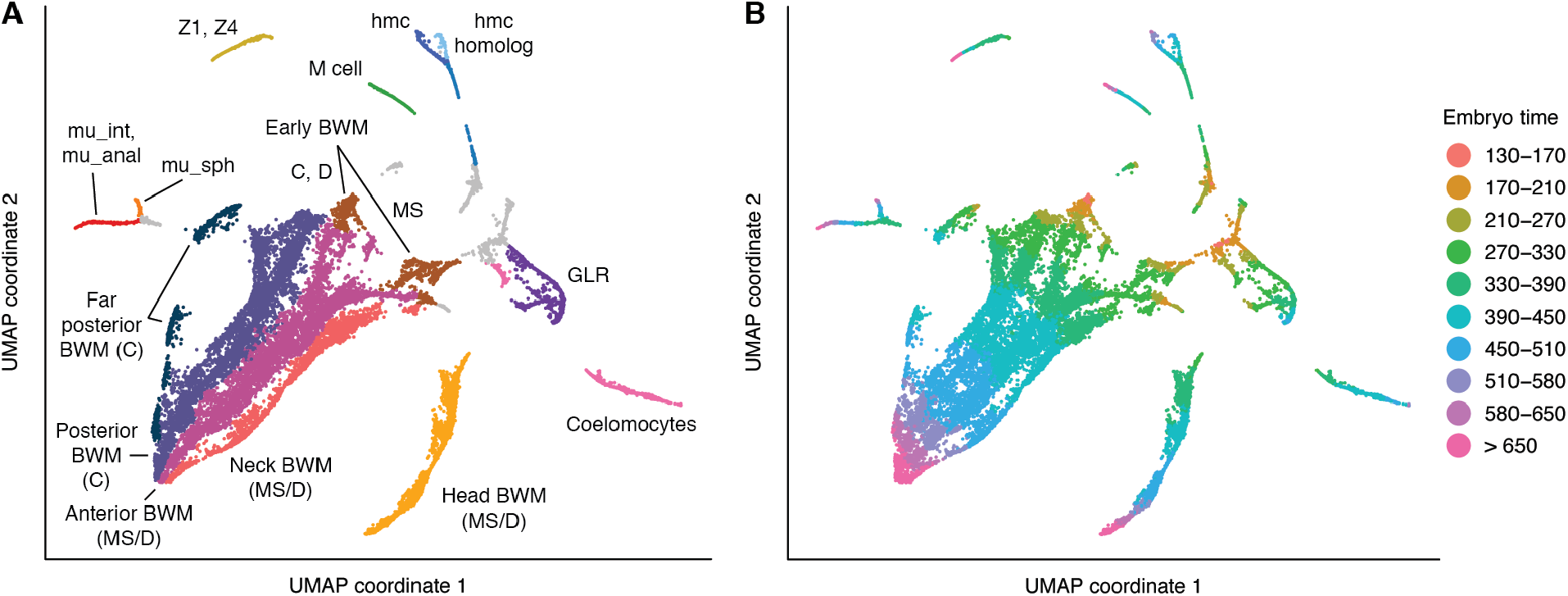
UMAP of body wall muscle and non-pharyngeal mesoderm cells. (**A**) Labels indicate cell types. See **Table S1** for marker genes used to identify cell types. MS, C, and D indicate cell lineages. Abbreviations: BWM = body wall muscle, mu_int = intestinal muscle, mu_anal = anal depressor muscle, mu_sph = anal sphincter muscle, hmc = head mesodermal cell. (**B**) Colors show estimated embryo times (minutes post first cleavage) for each cell.

**Fig. S6.**
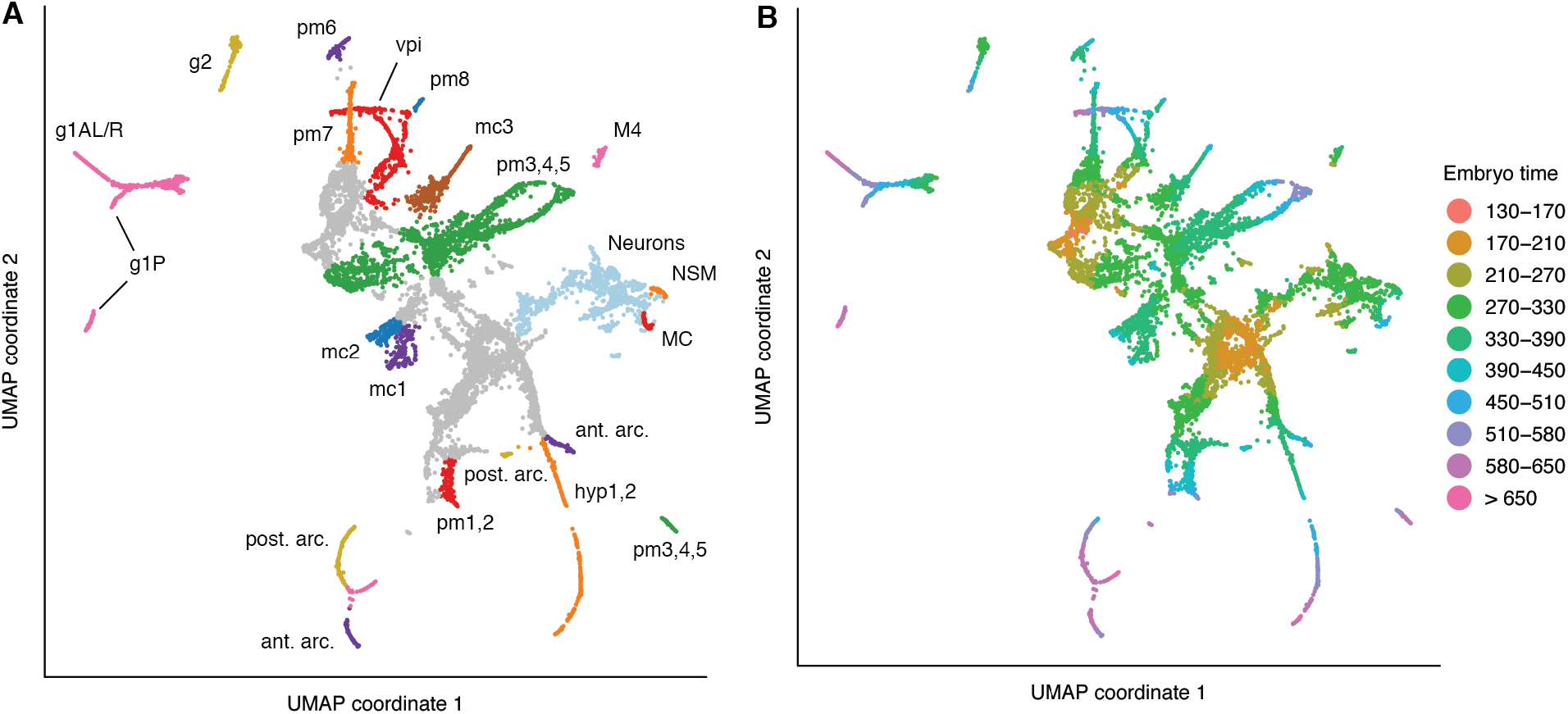
UMAP of pharyngeal cells. (**A**) Labels indicate cell types. See **Table S1** for marker genes used to identify cell types. Abbreviations: pm = pharyngeal muscle, mc = pharyngeal marginal cell, g1/g2 = pharyngeal gland, vpi = pharyngeal-intestinal valve, hyp = hypodermis, ant. arc. = anterior arcade cells, post. arc. = posterior arcade cells. Anterior and posterior arcades from late embryos converge in the UMAP to a common transcriptomic profile (pink cells at the bottom of the plot) (**B**) Colors show estimated embryo times (minutes post first cleavage) for each cell.

**Fig. S7.**
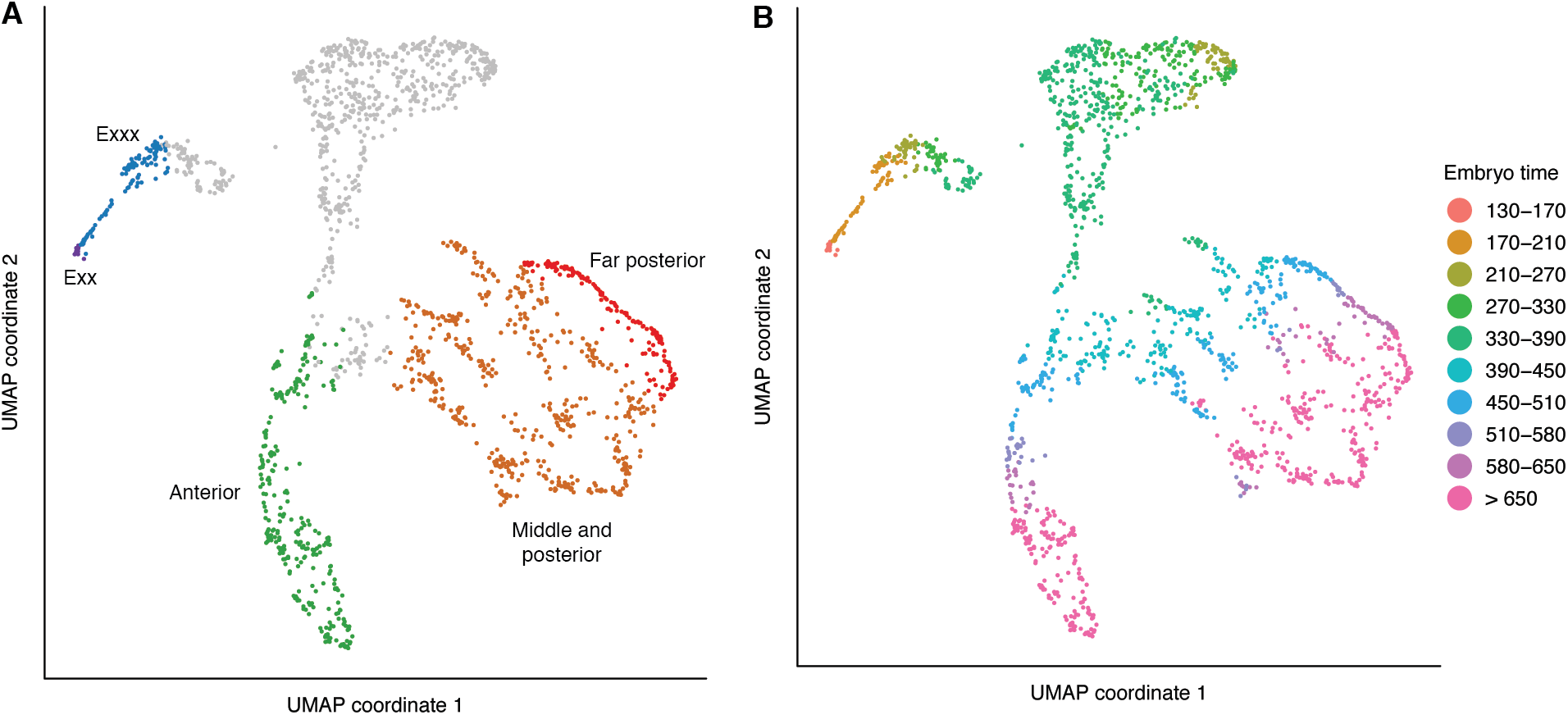
UMAP of intestine cells. (**A**) Labels indicate subsets of intestine cells and their relative position on the anterior-posterior axis. See **Table S1** for marker genes used to identify cell types. (**B**) Colors show estimated embryo times (minutes post first cleavage) for each cell.

**Fig. S8.**
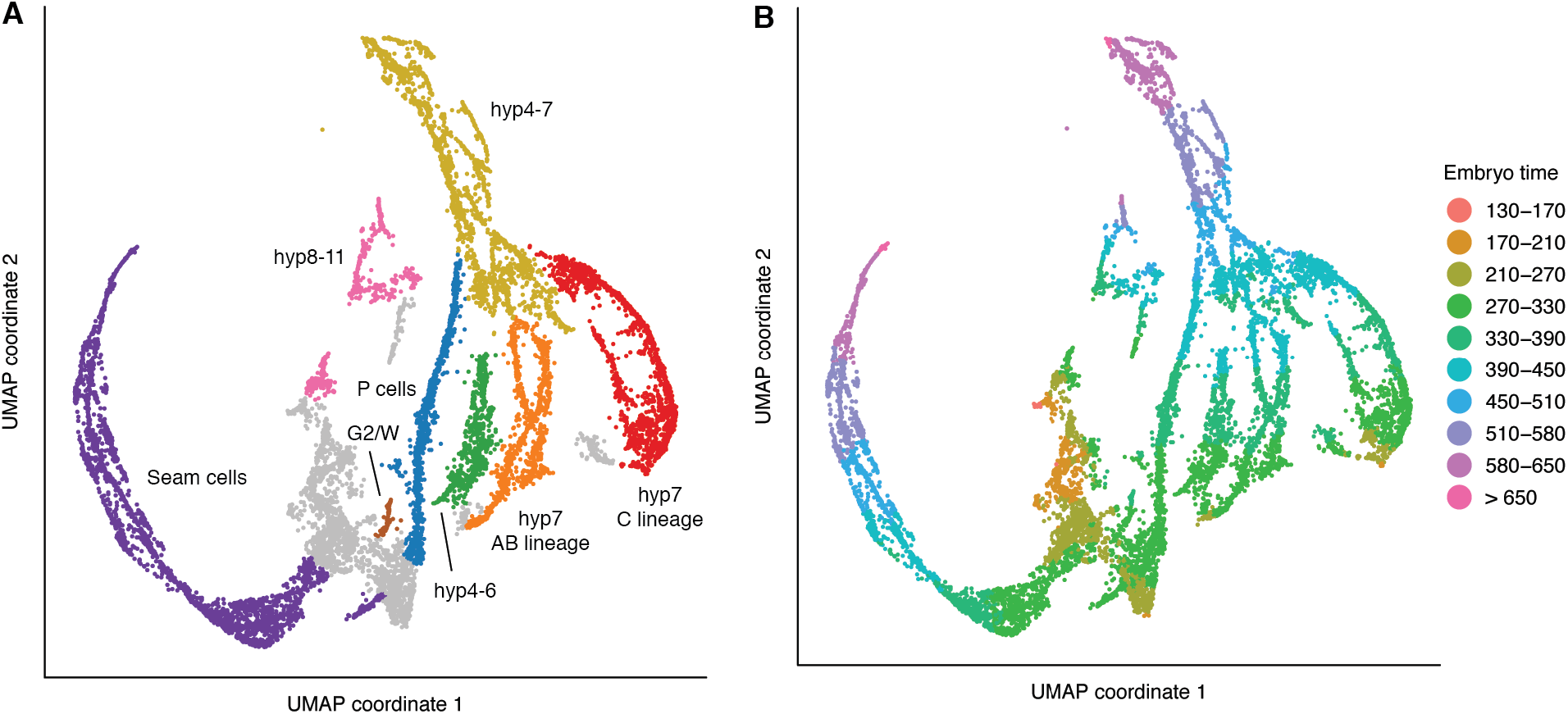
UMAP of hypodermis and seam cells. (**A**) Labels indicate cell types. See **Table S1** for marker genes used to identify cell types. hyp1-3 are not included in the UMAP. hyp1-2 appear in the pharynx UMAP (**Fig. S6**), and hyp3 appears in the glia UMAP (**Fig. S9**), consistent with their cell lineage (hyp1 and hyp2 are sisters/cousins of arcade cells, and hyp3 are sisters of ILsoDx). (**B**) Colors show estimated embryo times (minutes post first cleavage) for each cell.

**Fig. S9.**
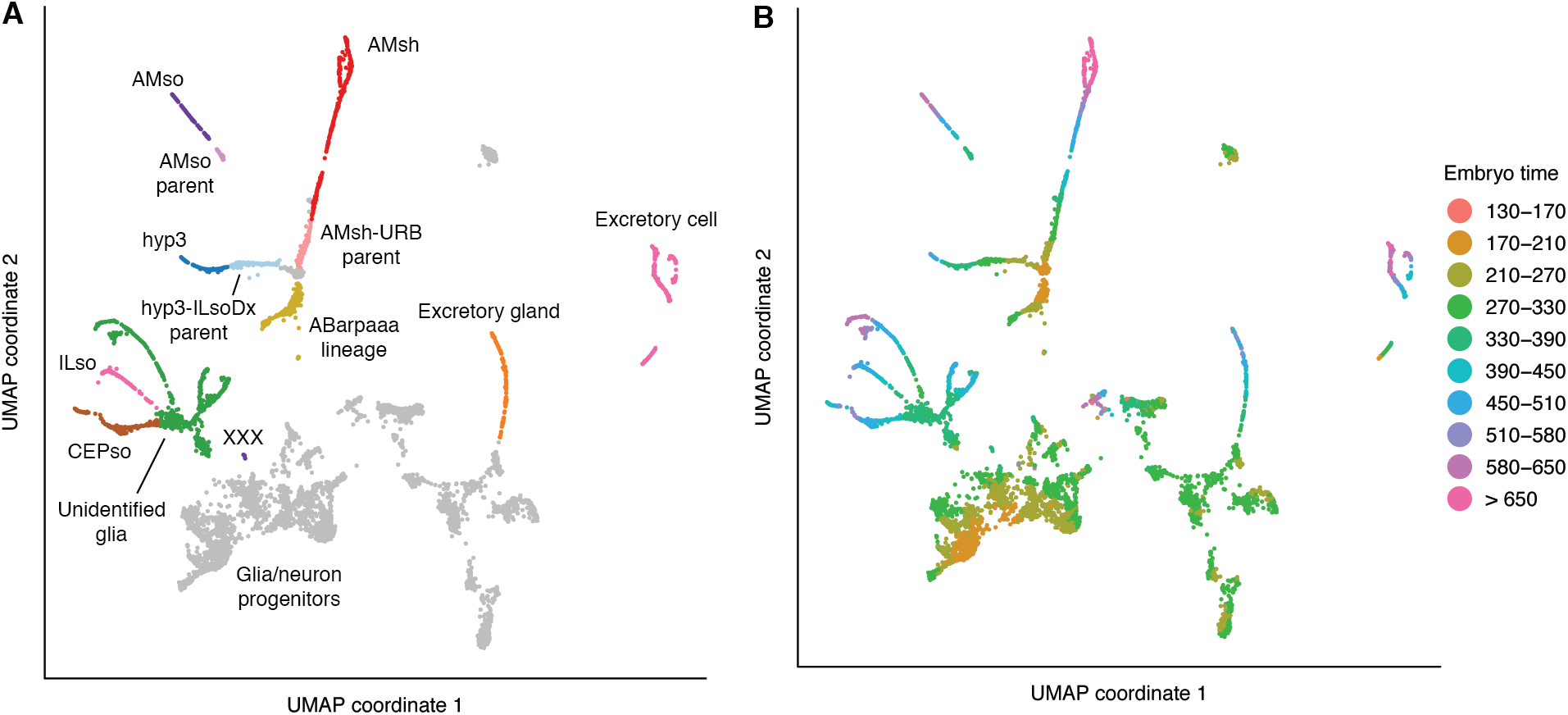
UMAP of glia, excretory cells, and progenitors. (**A**) Labels indicate cell types. See **Table S1** for marker genes used to identify cell types. See **Table S4** for marker genes used to identify pre-terminal cells. We have not yet been able to identify several glial cell types due to lack of suitable marker genes. Some non-glial/excretory cells, such as rectal cell progenitors, are also included in the UMAP. (**B**) colors show estimated embryo times (minutes post first cleavage) for each cell.

**Fig. S10.**
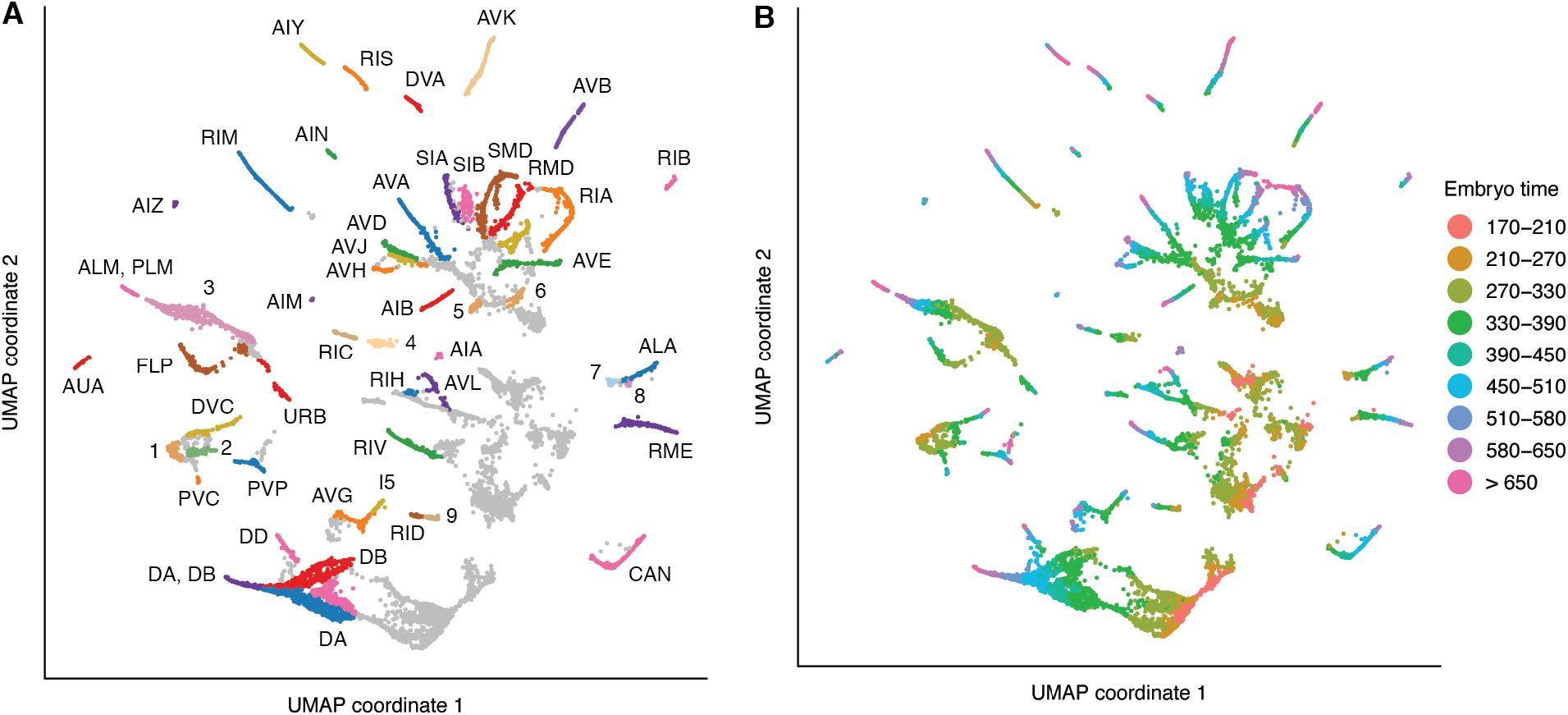
UMAP of non-ciliated neurons and progenitors. For a UMAP of ciliated neurons, see **Fig. 3A**. (**A**) Text labels indicate terminal cell types. See **Table S1** for marker genes used to identify cell types. Numeric labels indicate: **1** PHA-PVC-LUA neuroblast **2** PHB-HSN neuroblast **3** includes ALM-BDU and PLM-ALN neuroblasts, and possibly one or more of terminal ALM, PLM, ALN, and BDU before they become transcriptomically distinct **4** parent of RIC **5** parent of AVH **6** parent of RIA **7** ALA-RMED neuroblast **8** RMED, early after parent division **9** parent of RID. See **Table S4** for marker genes used to identify pre-terminal cells. (**B**) colors show estimated embryo times (minutes post first cleavage) for each cell.

**Fig. S11.**
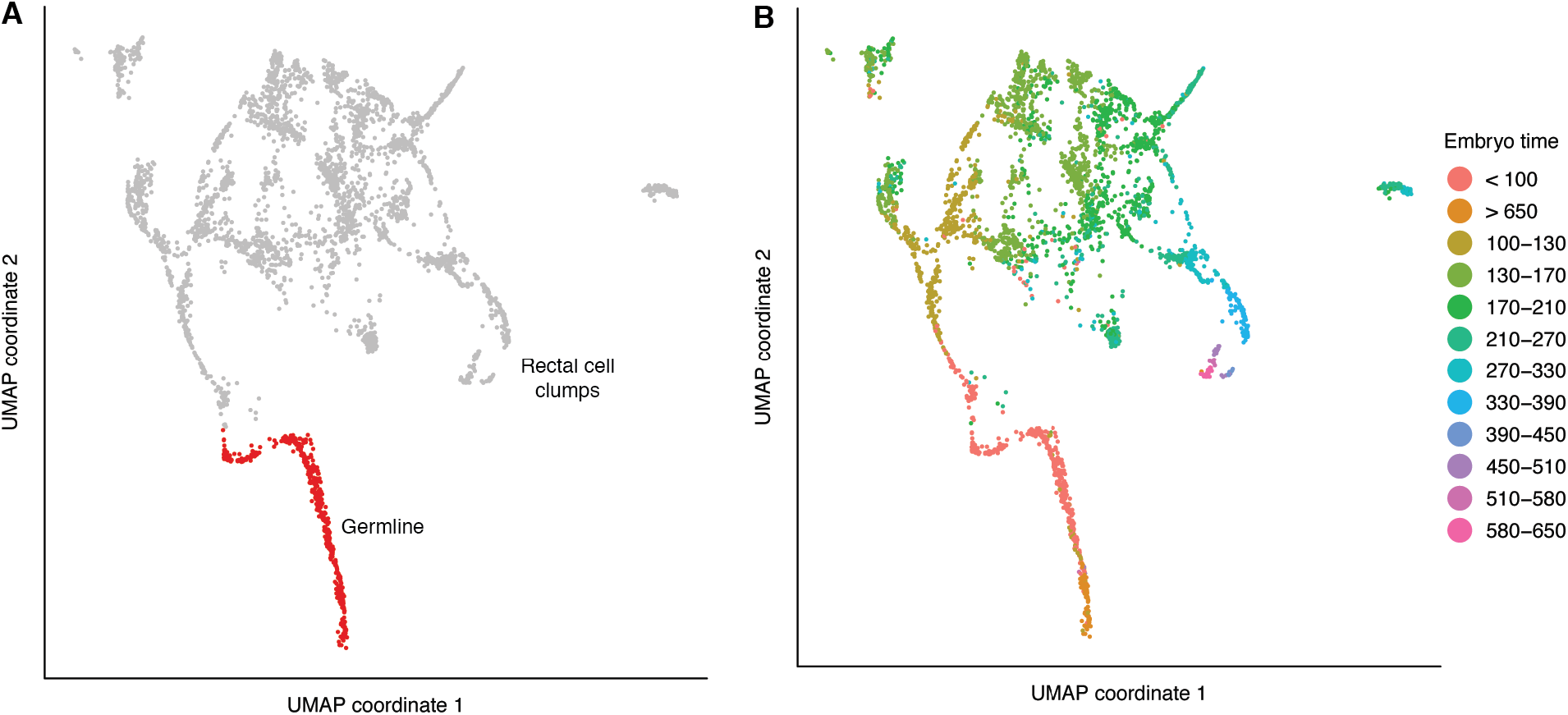
UMAP of early embryo, germline, and rectal progenitor cells. (**A**) Labels indicate cell types. The only noteworthy feature of this UMAP is the trajectory of germline development. See **Fig. S12** for time-specific UMAPs that better reveal the lineages of early embryo cells. Late stage rectal cells were either missing or appeared to be incompletely dissociated (i.e. cell clumps), so we did not attempt to annotate them in detail. (**B**) colors show estimated embryo times (minutes post first cleavage) for each cell. These estimates, which are based on correlation to a whole-embryo bulk RNA-seq time series, are inaccurate for germline cells, as genes that follow the same temporal dynamics for all somatic cells often have different expression dynamics in the germline, which is largely transcriptionally quiescent.

**Fig. S12.**
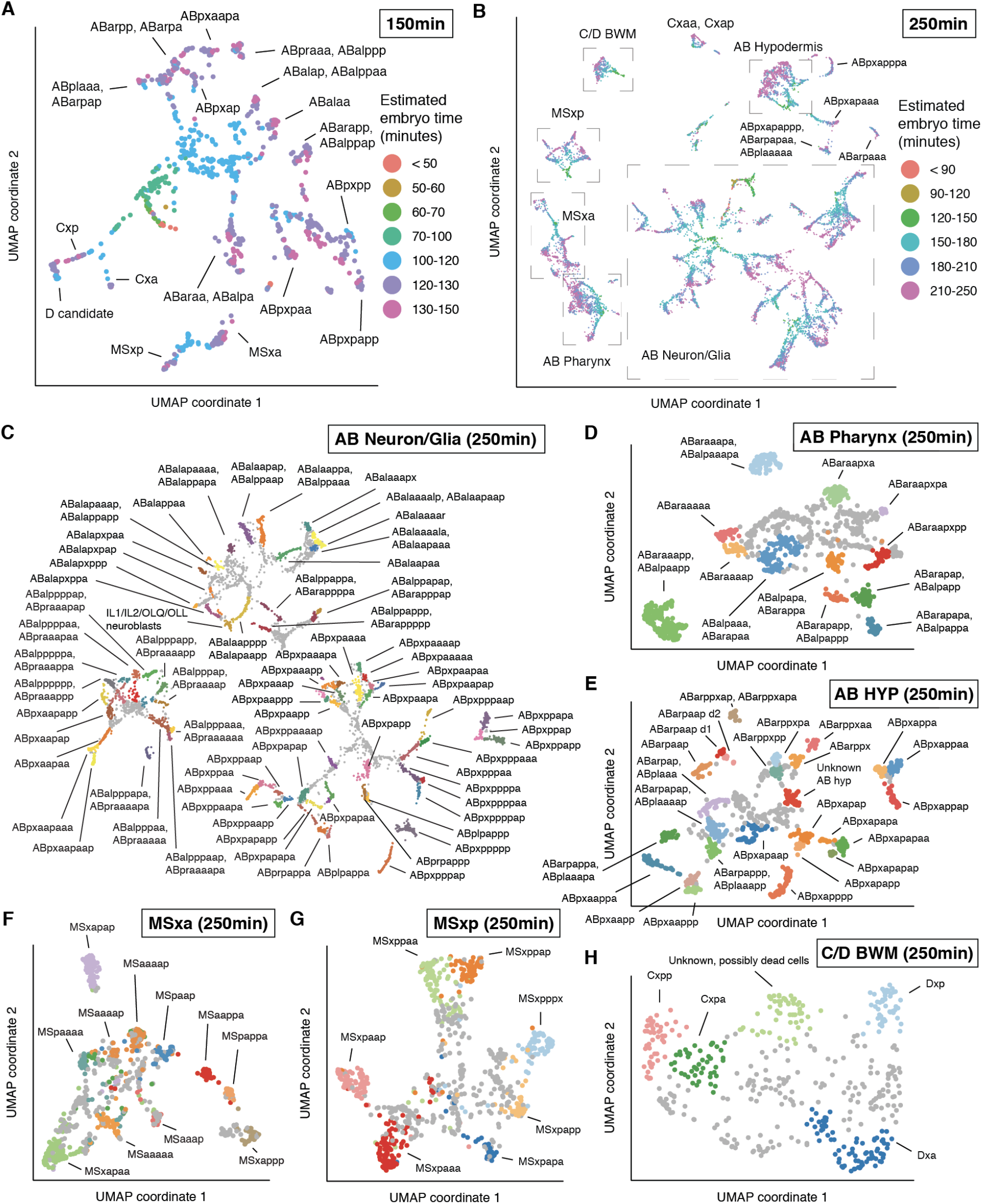
UMAP and detailed annotation of early cells. E lineage and germline cells are excluded from the UMAPs and were analyzed separately (See **Fig. S7, S11**). (**A**) Annotation of cells with estimated embryo time <= 150 minutes post first cleavage. (**B**) UMAP of early cells with estimated embryo time <= 250 minutes. This UMAP was further subdivided into (**C**) AB lineage cells that give rise to neurons and glia; (**D**) AB lineage cells that produce pharynx; (**E**) AB lineage cells that produce hypodermis; (**F**) the MSxa lineage; (**G**) the MSxp lineage and (**H**) C/D lineage cells that produce body wall muscle. The UMAPs in (**C-H**) are recomputed with variably expressed genes re-estimated with each individual subset of cells.

**Fig. S13.**
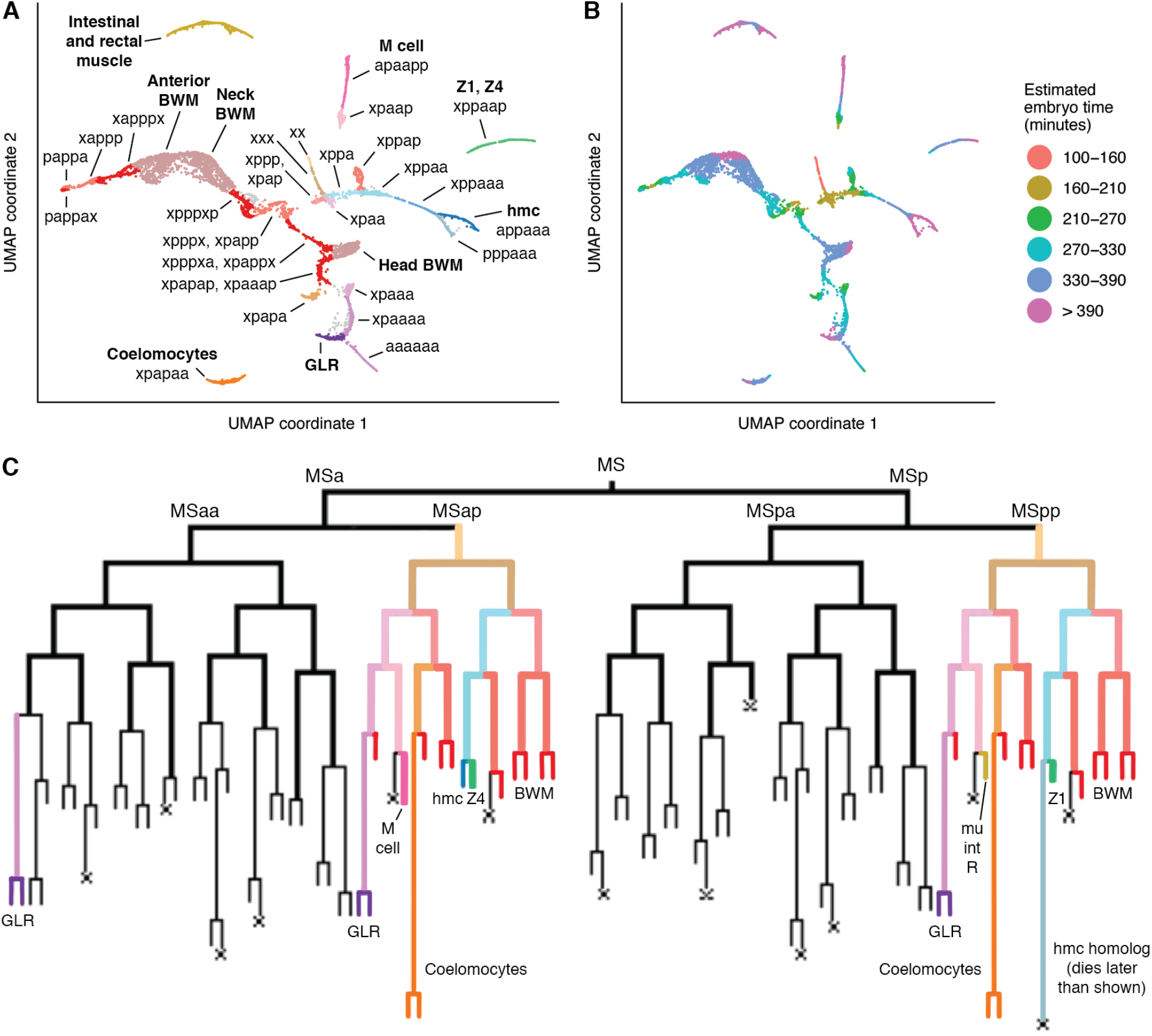
UMAP of non-pharyngeal mesoderm cells, focused on the early lineage. This UMAP includes the same cells as the muscle and mesoderm UMAP (**Fig. S5**), but excludes putative C and D lineage body wall muscle, MS lineage body wall muscle with estimated embryo time >400 minutes (post first cleavage), and coelomocytes with embryo time >400 minutes. (**A**) Text labels indicate MS lineages (i.e. “xppa” = MSxppa). Bold text labels indicate cell types. MSxppapx was not conclusively identified, but is presumed to be included in the head BWM cluster. (**B**) Estimated embryo time for each cell. (**C**) diagram of the MS lineage. Colored sub-lineages match the colors of cell groups in panel (**A**).

**Fig. S14.**
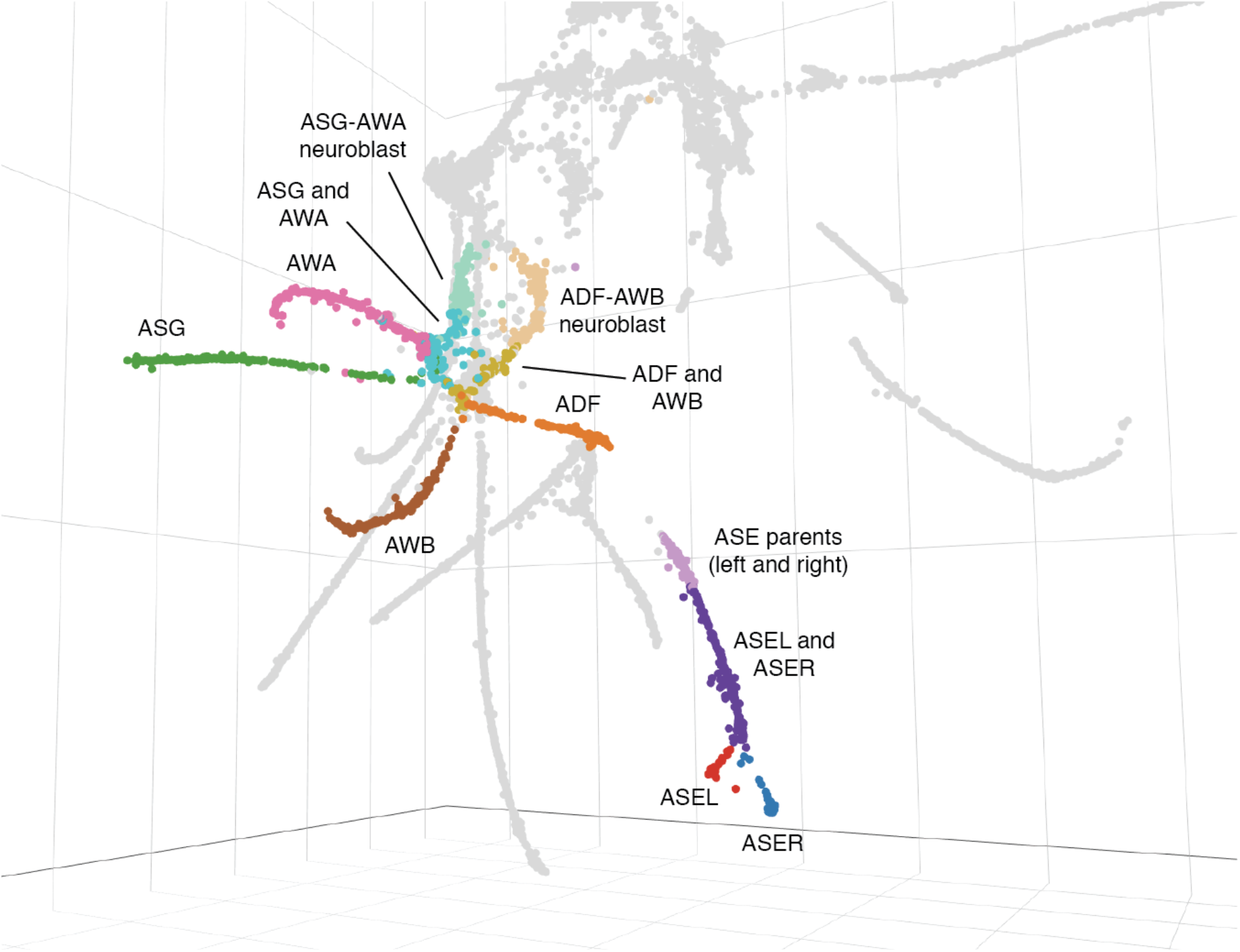
Ciliated neuron developmental trajectories are more continuous in a 3D UMAP. This plot is a screenshot of part of a 3D UMAP of ciliated neuron cells. The cells are the same as in **Fig. 3A**; the only difference is projecting into 3D instead of 2D. Developmental trajectories connecting the ASG-AWA and ADF-AWB neuroblasts to their respective daughter cells are continuous in this UMAP space, as is the branching trajectory of the left and right ASE neurons (ASEL and ASER). In the ASG-AWA and ADF-AWB trajectories, there are sections that appear before the branch points in the UMAP, but based on our embryo time estimates are likely to be terminal cells and not the parent neuroblasts. These sections may contain both daughter cells of each trajectory after their birth but before they differentiate. Cells in the “ADF and AWB” section co-express in the same cells the marker genes *lag-1*, which persists only in ADF, and *lim-4*, which persists only in AWB; however their estimated embryo times span ∼100 minutes after the parent cells’ division time. Note that the grey, unannotated cells below the ADF trajectory are behind the ADF cells in 3D space, as are the grey cells overlapping the AWB trajectory.

**Fig. S15.**
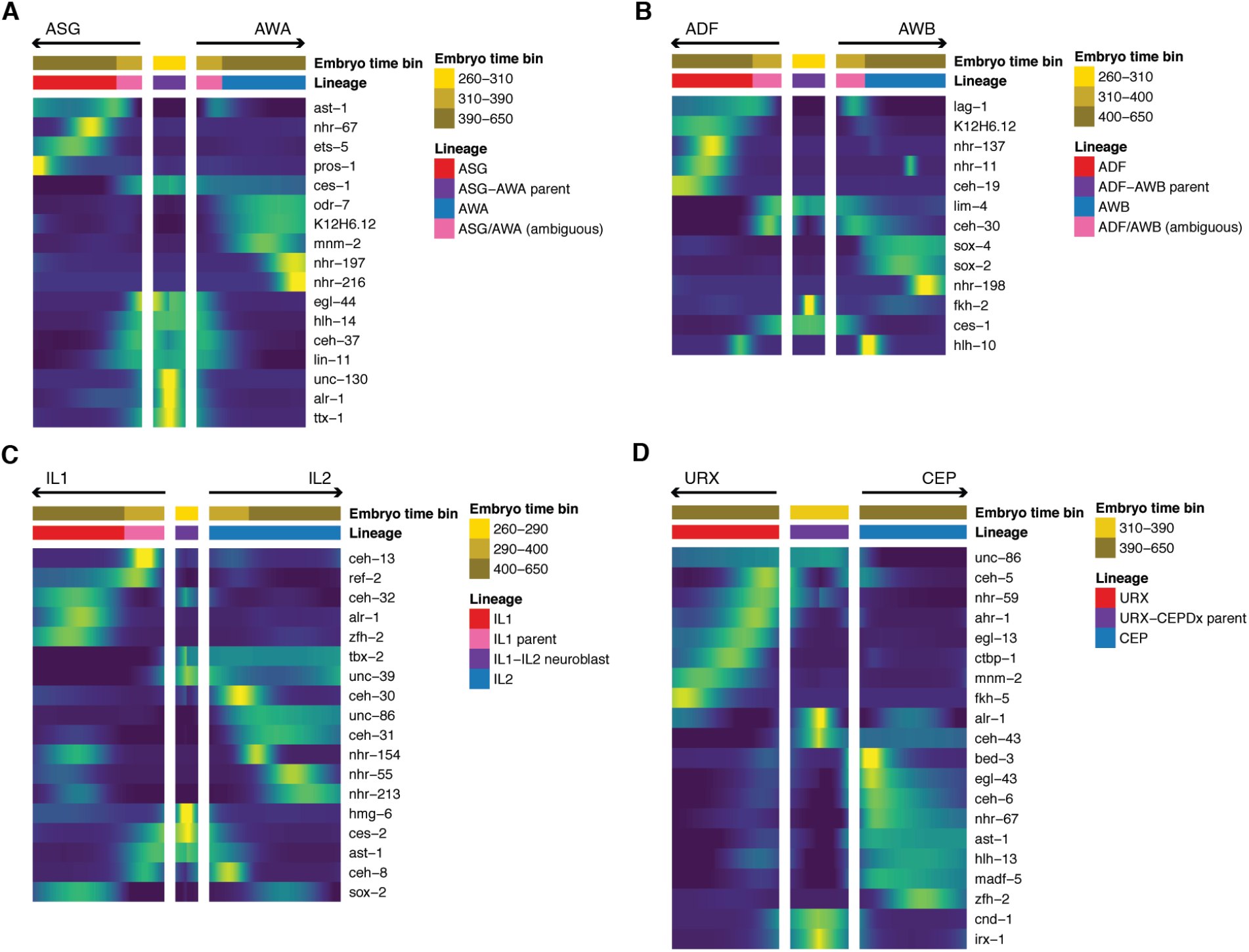
Differentially expressed transcription factors associated with ciliated neuron lineage branches. Heatmaps showing patterns of differential transcription factor expression associated with branches in (**A**) the ASG-AWA lineage, (**B**) the ADF-AWB lineage, (**C**) the IL1-IL2 lineage, and (**D**) the URX-CEPDx lineage. A heatmap for the ASE-ASJ-AUA lineage is shown in **Fig. 3D**. Expression values are log-transformed, then centered and scaled by standard deviation for each row (gene). In each of the ASG-AWA and ADF-AWB lineages, there is a set of cells that are before the branch point of the trajectory in UMAP space (see **Fig. S14**), but based on embryo time estimates and marker gene expression patterns, are likely to be terminal cells. In the ADF-AWB lineage, these cells co-express *lag-1*, which is selectively retained in ADF, and *lim-4*, which is selectively retained in AWB, suggesting that this cell set may include undifferentiated, terminal ADF and AWB cells.

**Fig. S16.**
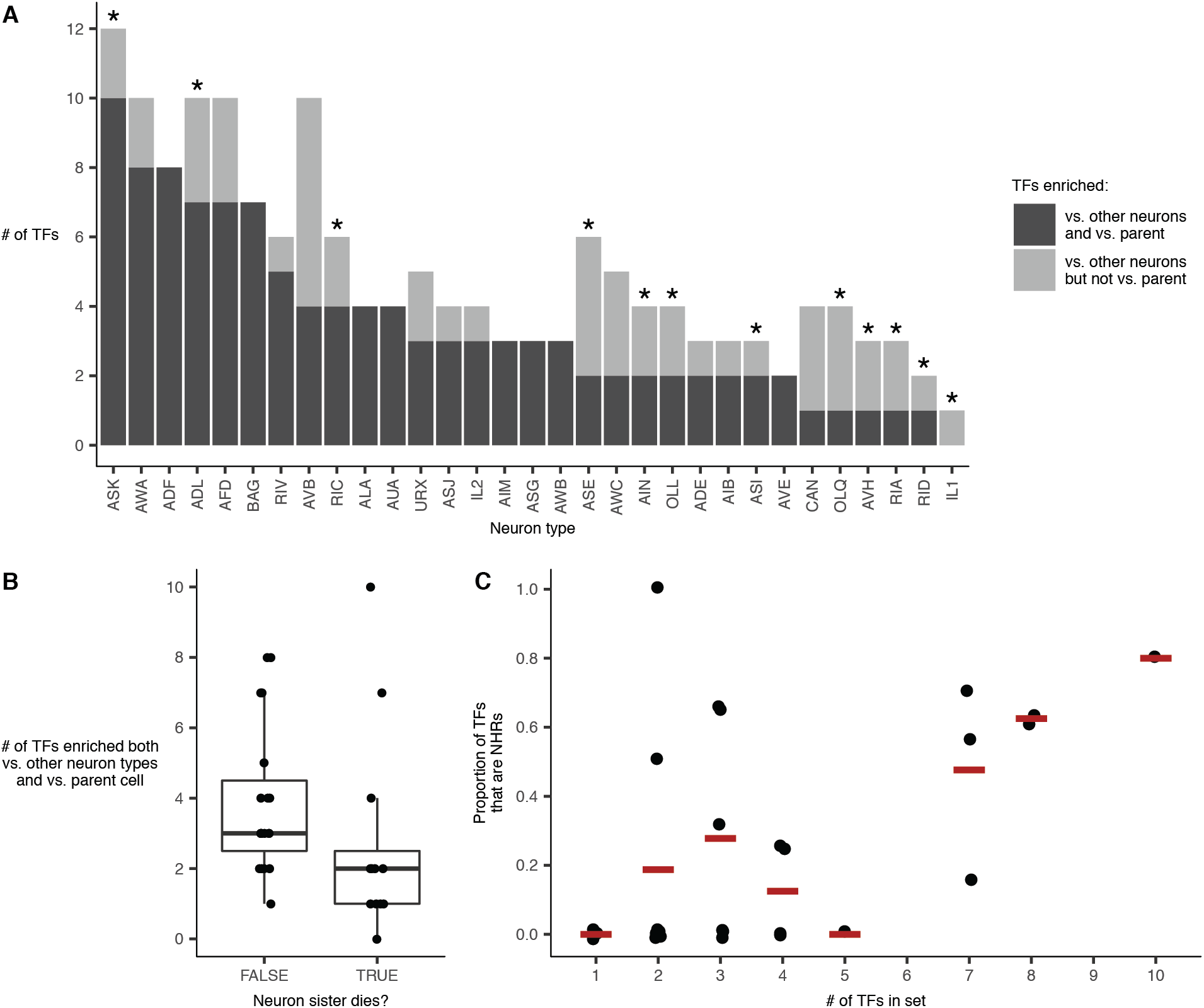
Statistics related to differential expression of TFs in terminal neurons. Plots in this figure consider the set of TFs that are robustly expressed in a neuron type (>100 TPM), enriched in the neuron type vs. all other neurons (>8-fold), and enriched in the terminal neuron vs. its parent (>8-fold). (**A**) shows the number of TFs meeting these criteria for each neuron type in which we annotated both the terminal neuron and its parent. Asterisks indicate lineages in which the parent neuroblast produces one neuron and one cell that dies. (**B**) compares the distribution of these TF counts for lineages in which one daughter cell dies vs. lineages in which both survive. The difference in distributions is marginally significant (p = 0.032, Wilcoxon rank sum test). (**C**) shows, for each neuron type (points), the proportion of the set of TFs that meet the criteria that are nuclear hormone receptors (*nhr-* genes). Outlier neuron types that have a high number of TFs enriched both vs. other neurons and the neuron’s parent express several cell type specific *nhr-* genes.

**Fig. S17.**
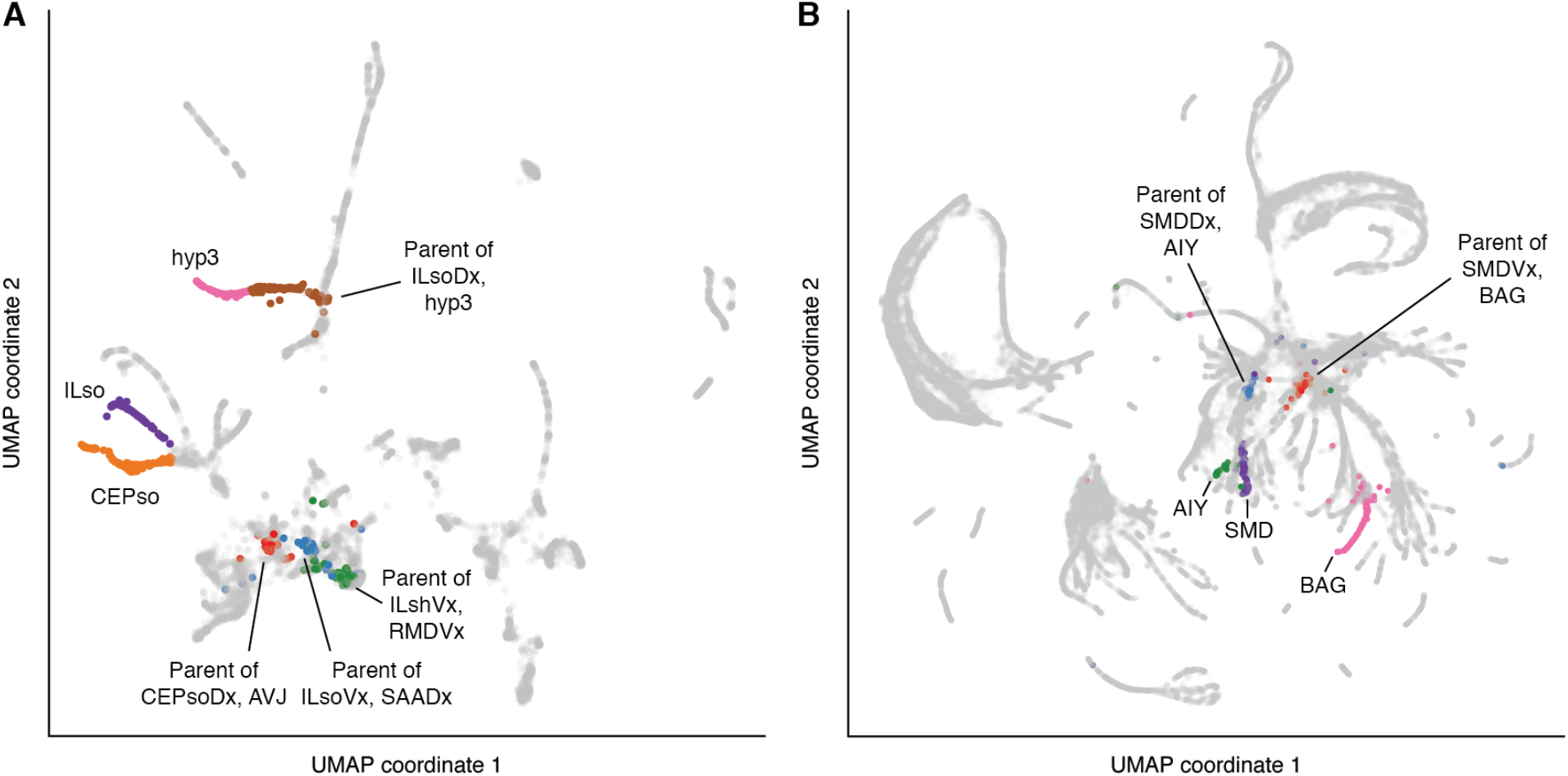
Examples of lineages that form discontinuous trajectories in UMAP space. (**A**) Plot is a UMAP of glial cells and progenitors (same as **Fig. S8**). Labels show selected cell type and lineage annotations. Other cells in the cluster with CEPso and ILso are also glia, but we were unable to narrow them down to specific cell subtypes. Parents of CEPsoVx and ILsoL/R were not identified. (**B**) Plot is a UMAP of all cells in the dataset (same as **Fig. S3**). Labels show cells in two lineages that produce SMD neurons. See **Tables S1** and **S4** for marker genes used in cell type and lineage annotations. Pre-terminal cells in both panels were annotated based on the UMAPs in **Fig. S12**, not the UMAPs shown.

**Fig. S18.**
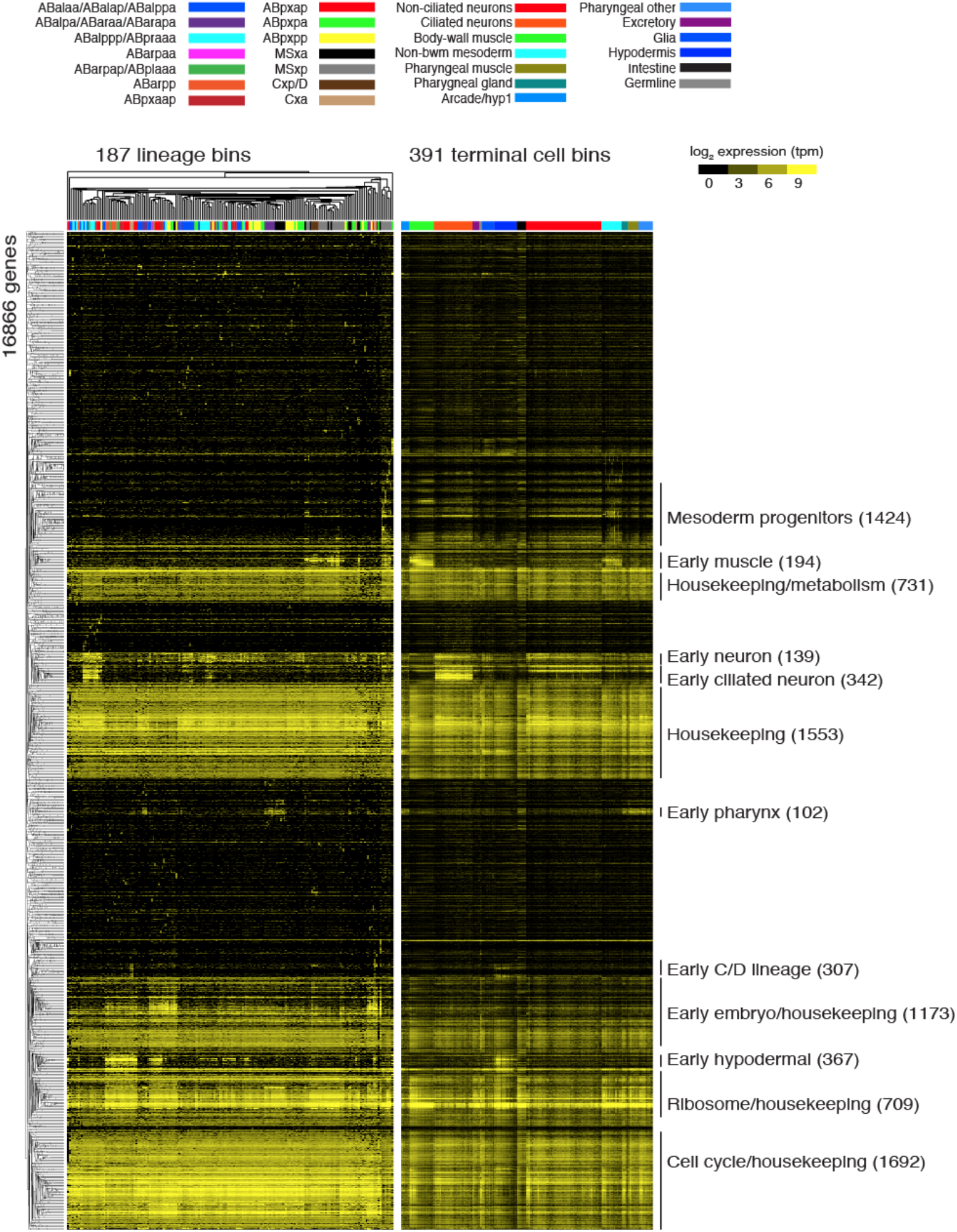
Hierarchical clustering of early lineage bins. This heatmap shows the log_2_ expression of all genes (rows) that are expressed (TPM > 0) in at least one early lineage bin (columns). Genes and lineage bins are ordered by hierarchical clustering of the early lineage data. The right panel shows the expression values in terminal cell bins, with genes (rows) ordered by the clustering as generated from the early lineage cell bins and terminal cell bins (columns) ordered as in **Fig. S19**.

**Fig. S19.**
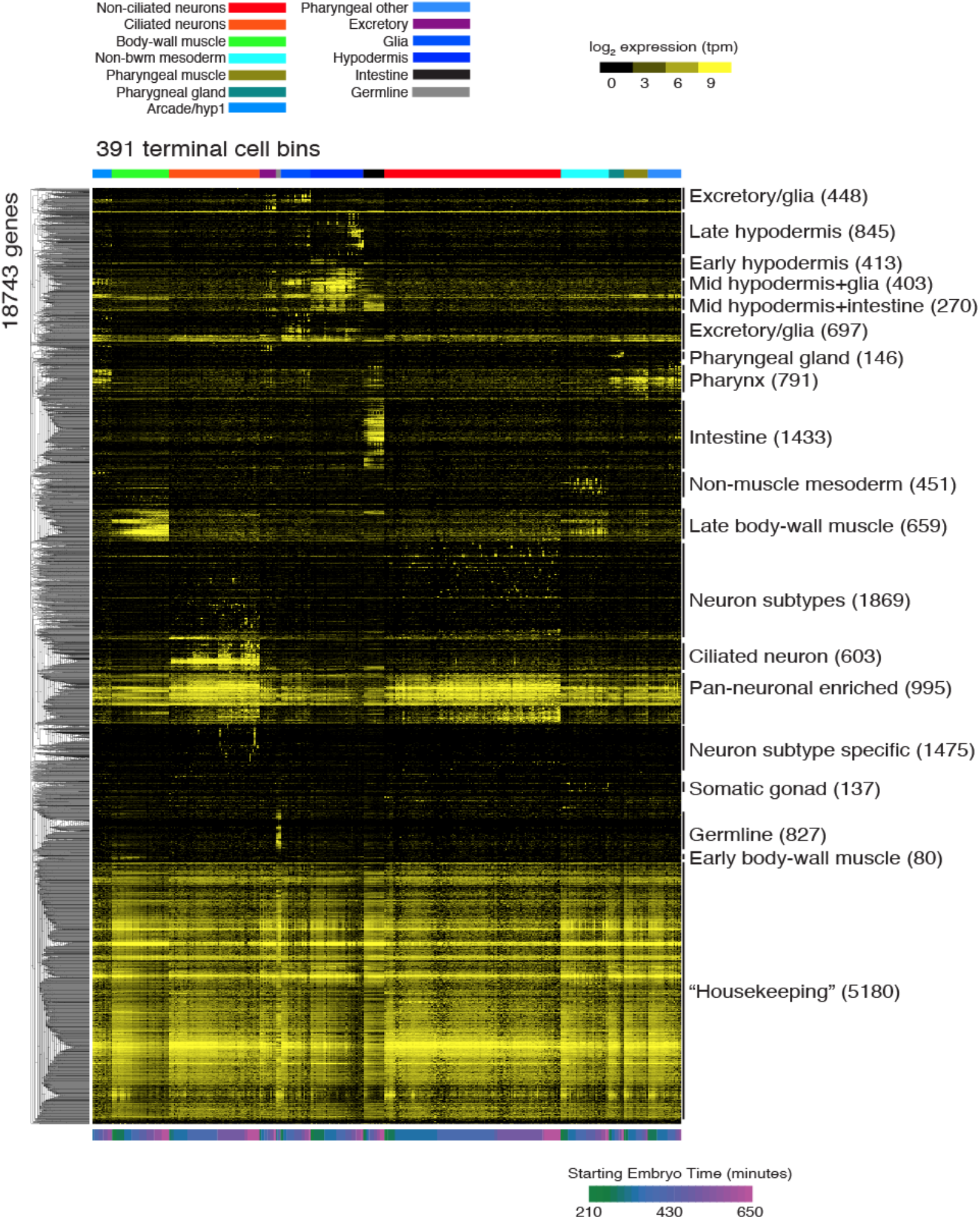
Hierarchical clustering identifies signatures of tissue and cell type differentiation. This heatmap shows the log_2_ expression (median bootstrap tpm plus a pseudocount of 1) of all genes (rows) that are expressed (TPM > 0) in at least one terminal cell bin (columns). Genes are ordered by hierarchical clustering, and cell bins are ordered by tissues (colored as in the legend), and within tissues by the beginning of the time bin in minutes (early to late). Gene clusters are labeled by sites of predominant expression. Numbers in parentheses are the number of genes in that cluster.

**Fig. S20.**
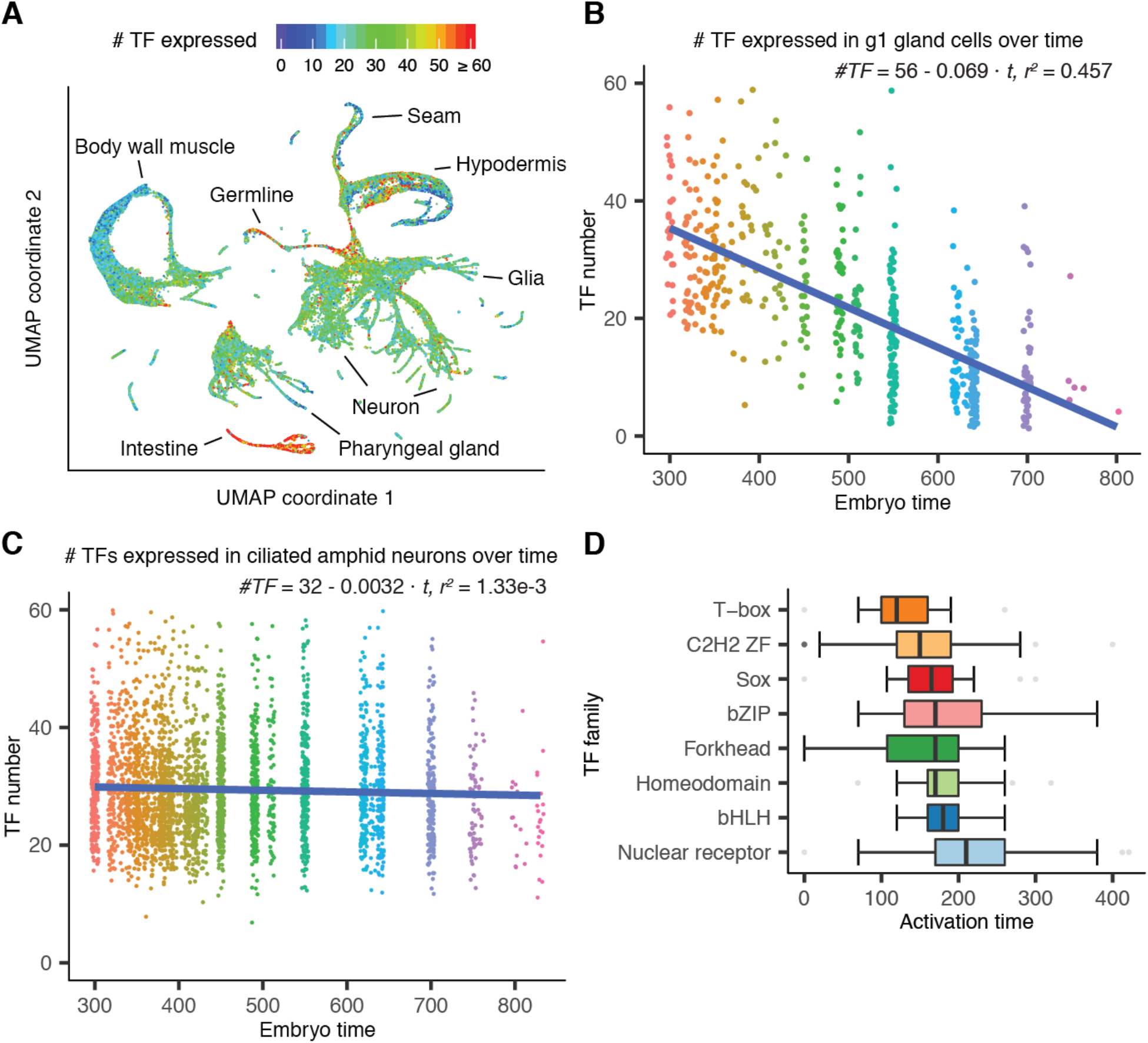
Global expression patterns of transcription factors. (**A**) Global UMAP colored by number of expressed TFs in each cell, with max value cut off at 60 (∼95th percentile). (**B**) Number of TF expressed in g1 gland over time. Equation shows linear regression result. Points are colored by estimated embryo time. (**C**) Number of TF expressed in ciliated amphid neurons over time. Similar to C, the data were fit using a linear model. (**D**) Box plot showing TF activation times—the embryo time when a TF first becomes expressed—grouped by TF family. For each TF, its activation time is defined as the 5th percentile of the estimated embryo time values for cells that express that TF. TF families that have fewer than 10 members detected in the current dataset were excluded from this plot.

**Fig. S21.**
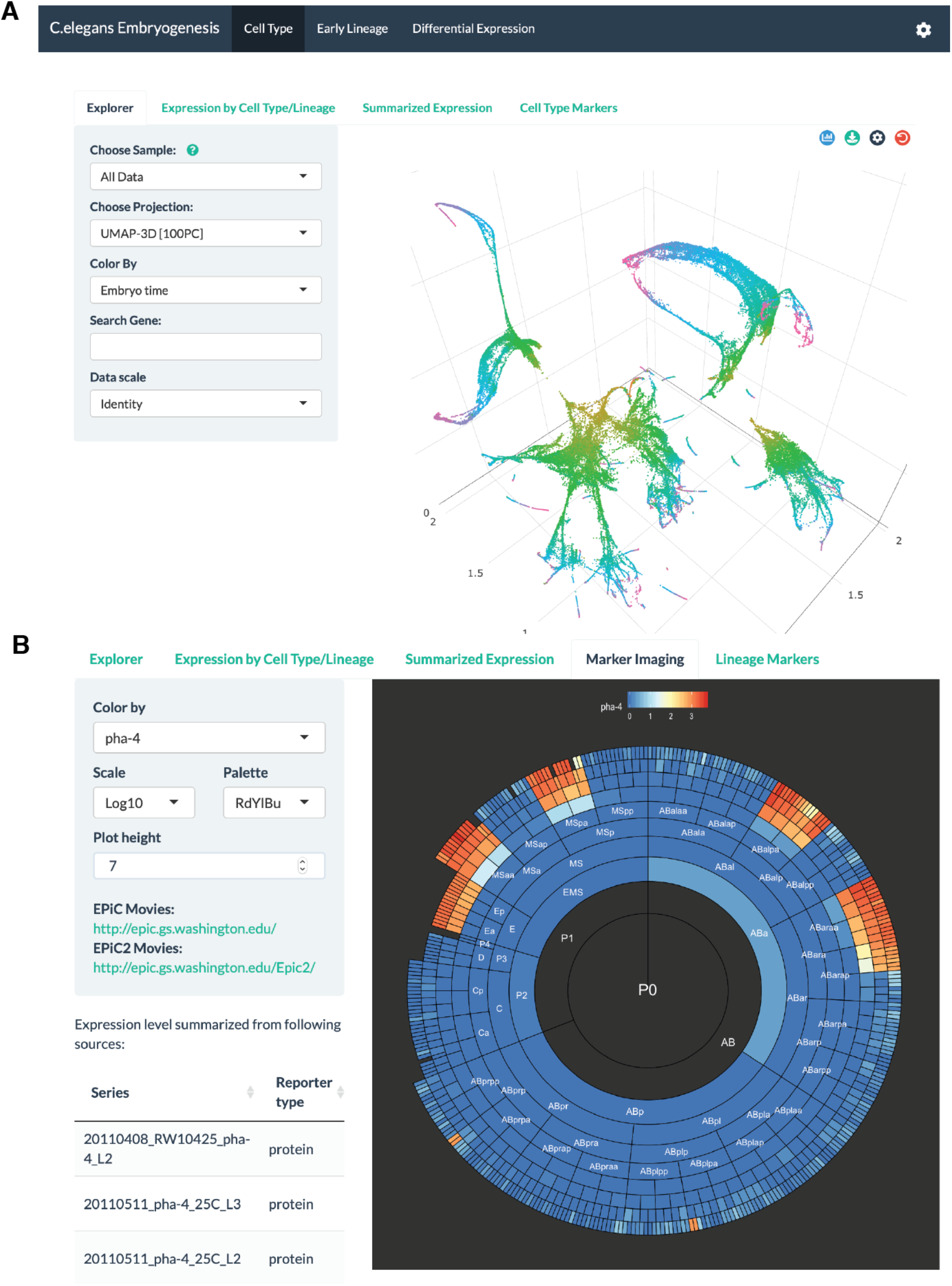
Screenshots of VisCello. (**A**) Screenshot of the cell type explorer, which enables interactive visualization of 2D and 3D UMAPs and PCA plots for different subsets of the data. The view shown in the panel is a 3D UMAP for all cells colored by estimated embryo time. Users can overlay gene expression, cell type, number of expressed genes and other statistics on this plot. The cell type explorer also features box/violin plot for gene expression across cell types, lineages or time, summarized gene expression tables, and marker gene tables. (**B**) Screenshot of the early cell lineage explorer, which enables interactive visualization and comparison of the sc-RNA-seq data and summarized live imaging data. Panel shows a radiograph of average fluorescent intensity (log10 scaled) of *pha-4*, measured by live imaging.

**Fig. S22.**
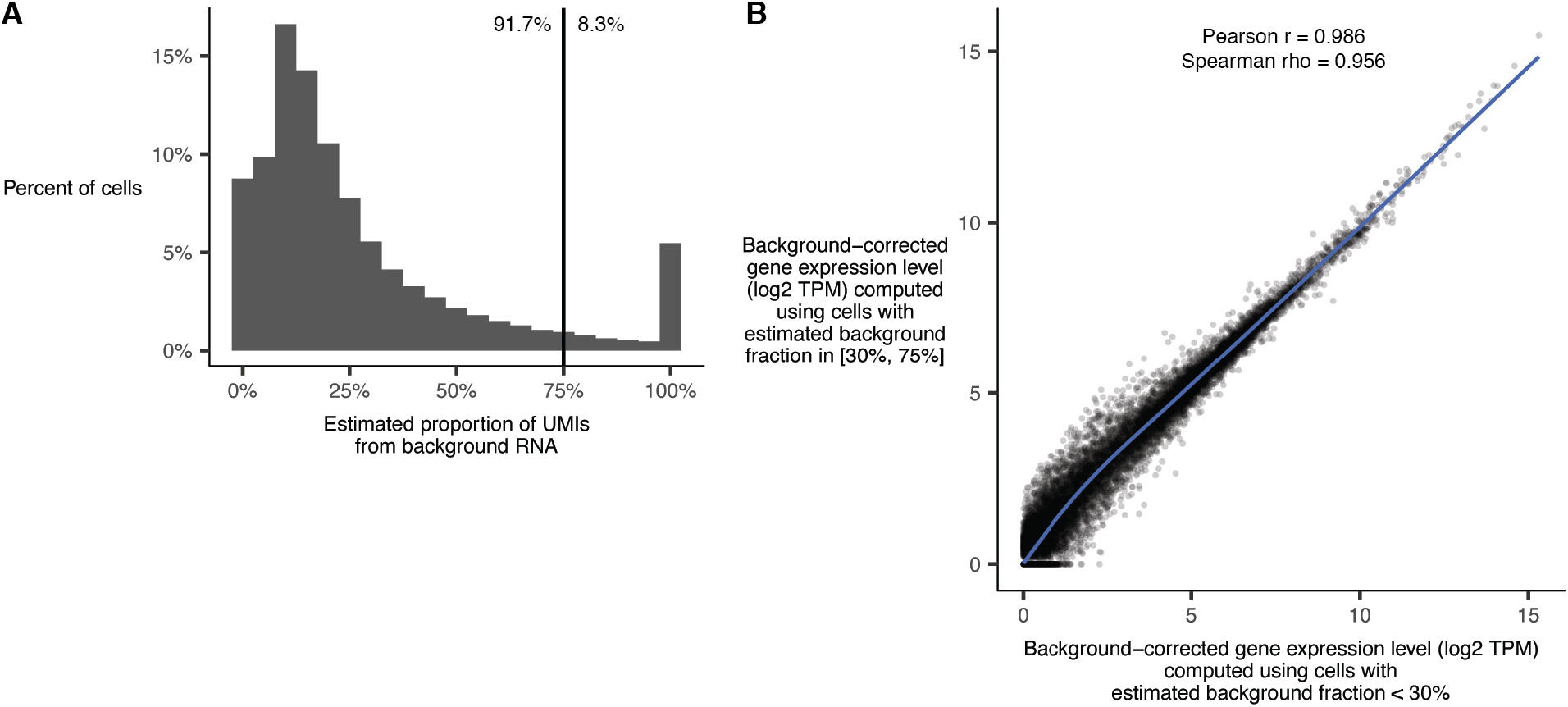
Distribution of estimates for the proportion of UMIs in a cell that come from background RNA. (**A**) The process for making the estimates is described in the methods section “Per-cell background correction and filtering”. Due to the sparsity of the single cell data, the estimates are noisy. Numbers to the left and right of the vertical line indicate the proportion of cells with estimated background fraction < or >= 75%. Cells with background fraction >= 75% are filtered from all downstream analyses. (**B**) After per-cell background correction, cells with low and high background fractions have near-identical average gene expression profiles. Plot shows average gene expression profiles (measured in transcripts per million) computed from non-head body wall muscle cells divided into two groups: cells with estimated background fraction < 30% (x axis) and cells with background fraction in the range [30%, 75%].

